# Artesunate, EDTA and colistin work synergistically against MCR-negative and -positive colistin-resistant *Salmonella*

**DOI:** 10.1101/2024.05.07.593013

**Authors:** Yajun Zhai, Peiyi Liu, Xueqin Hu, Changjian Fan, Xiaodie Cui, Qibiao He, Dandan He, Xiaoyuan Ma, Gongzheng Hu, Yajun Zhai

## Abstract

Discovering new strategies to combat the multi-drug resistance bacteria constitutes a major medical challenge of our time. Previously, artesunate (AS) has been reported to exert antibacterial enhancement activity in combination with β-lactam antibiotics, via inhibition of the efflux pump AcrB. However, combination of AS and colistin (COL) revealed weak synergistic effect against a limited number of strains, and few studies have further explored its possible mechanism of synergistic action. In this paper, we found that AS and EDTA could strikingly enhance the antibacterial effects of COL against *mcr*-*1*^-^ and *mcr*-*1*^+^ *Salmonella* strains either *in vitro* or *in vivo*, when used in triple combination. The excellent bacteriostatic effect was primarily related to the increased cell membrane damage, accumulation of toxic compounds and inhibition of MCR-1. The potential binding sites of AS to MCR-1 (THR283, SER284, and TYR287) were critical for its inhibition of MCR-1 activity. Additionally, we also demonstrated that the CheA of chemosensory system and virulence-related protein SpvD were critical for the bacteriostatic synergistic effects of the triple combination. Selectively targeting CheA, SpvD or MCR using the natural compound AS could be further investigated as an attractive strategy for treatment of *Salmonella* infection. Collectively, our work opens new avenues towards the potentiation of COL and revealed an alternative drug combination strategy to overcome COL resistant bacterial infections.

## Introduction

*Salmonella* is globally recognized as a major zoonotic foodborne pathogen, that responsible for food poisoning, gastroenteritis and even life threatening in animals and humans (Galán-Relaño *et al*, 2023, Gangathraprabhu, 2020). Antibiotics are commonly used to shorten duration of illness and reduce infectivity. However, with the increasing use of antibiotics, the infections caused by multi-drug resistant (MDR) pathogens, especially the carbapenemase-producing *Enterobacteriaceae*, have become the major source of public-health concerns (Foletto *et al*, 2021). The shortage of new antibiotics for these MDR bacteria strains has led to the re-use of polymyxins as the “last resort” antimicrobial drug with the inevitable risk of emerging resistance (Falagas & Kasiakou, 2005). Colistin (polymyxin E, COL), a fatty acyl oligopeptide antibiotic, is an active agent against Gram-negative (G^-^) pathogens, and has been widely used to combat *Salmonella* infections. Generally, COL kills bacteria through a detergent-like effect, that the polycationic ring of COL electrostatically interacted with the cell envelope components, causing the competitive displacement of divalent cations calcium (Ca^2+^) and magnesium (Mg^2+^), destabilizing the membrane, thus killing the bacterium via the “self-promoted uptake” pathway (Kaye *et al*, 2016). Beyond that, other models for the antibacterial activity have been reported, including vesicle-vesicle contact, hydroxyl radical death, inhibition of respiratory enzymes and anti-endotoxin COL activity pathways (El-Sayed Ahmed *et al*, 2020). Until now, numerous chromosomally- or plasmid-mediated mechanisms underlying polymyxins resistance in G^-^ bacteria have been identified, including intrinsic, mutation (eg. PmrAB, PhoPQ or AcrAB-TolC mutants), adaptation mechanisms or horizontally acquired resistance via the phosphoethanolamine (pEtN) transferase genes *mcr-1* to *9* (Carroll *et al*, 2019; Lima *et al*, 2018; Poirel *et al*, 2017).

To tackle the increasing emergence of MDR pathogens, many alternative therapies, less costly and time-consuming than drug discovery, have been new areas of current research interest, involving the combination therapy of existing agents, the drug discovery from natural products, and the evaluation of drug resistance reversers (Rosenthal, 2003). A variety of antibiotic adjuvants that may or may not have direct antibacterial effects have been widely investigated to increase the effectiveness of current antibiotics or delay the emergence of drug resistance, such as β -lactamase inhibitors, aminoglycoside-modifying enzyme inhibitors, membrane permeabilisers, and efflux pump inhibitors (Laws *et al*, 2019). Several promising inhibitors, for example zidebactam and pyrazolopyrimidine compounds have been described as the β-lactam and aminoglycoside enhancers against G^-^ bacteria (Moya *et al*, 2017; Stogios *et al*, 2013). Alternatively, numerous synthetic antimicrobial peptides and plant-derived natural products have been shown to possess membrane permeabilising or efflux pump inhibitory activity, with the combination of azithromycin, ciprofloxacin, imipenem *et al* (Aron & Opperman, 2016; Lin *et al*, 2015; Su & Wang, 2018).

Artesunate (AS) is a semi-synthetic derivative of anti-malarial compound artemisinin that extracted from the traditional Chinese herb Artemisia annua. Beyond remarkable antimalarial action, AS and other artemisinin derivatives, eg. dihydroartemisinin (DHA), have been proven to restored the antibacterial effect of COL against *Escherichia coli* (*E. coli*), while themselves didn’t exhibited intrinsic antimicrobial activity against clinical *E. coli* isolates as well as ATCC 25922 (Wei *et al*, 2020; Zhou *et al*, 2022). In addition, AS has also been proved to enhance the effectiveness of various β-lactam and fluoroquinolones antibiotics against MDR *E. coli* via inhibiting the efflux pump AcrAB-TolC (Pan *et al*, 2020; Wei *et al*, 2020). Nonetheless, the synergistic effect between AS and COL were only observed in a limited number of strains, with a modest reversal effect. Therefore, few studies have been undertaken to evaluate its underlying mechanism. Under this circumstance, it is meaningful to explore new drug combinations between AS and COL against MDR bacteria. Encouragingly, in this study, we confirmed the prominent synergistic effects of AS and EDTA to restore the antimicrobial activity of COL and its possible molecular mechanism.

## Results

### Artesunate and EDTA could enhance the effects of colistin against *Salmonella* strains

The antimicrobial activities of AS, EDTA or COL alone were initially investigated to the COL-sensitive strains of *Salmonella* (JS, S34), COL-resistant clinical strains of *Salmonella* (S16, S20, S13, and S30), *E*. *coli* (E16), and intrinsically COL-resistant species (*Morganella morganii* strain M15, *Proteus mirabilis* strain P01). Results showed that AS or EDTA alone had no direct antibacterial activity against these strains, with the minimum inhibitory concentrations (MICs) 1250 or > 125 mg/L (Table supplement 1). Excepting for the two intrinsically COL-resistant strains M15 and P01, there were only slight decrease of COL MICs for other strains (fold changes ranging from 0 to 133), when the sub-inhibitory concentrations (1/4, 1/8, 1/16 MIC) of AS or EDTA were combined with COL (namely AC or EC) (Table supplement 1). Whease, we found marked decrease of COL MICs (up to 60000 fold), after three drug combinations (namely AEC) (Table 1). These results indicated that when used simultaneously with COL, AS and EDTA exerted antibacterial enhancement activity for *Salmonella* and *E*. *coli*, but not the intrinsically COL-resistant species. Thus, AS and EDTA could be considered as adjuvants to reverse COL resistance in *Salmonella*.

**Table 1.** The antibacterial activities of COL against the tested strains after single and triple combinations.

| Strains | *mcr-1* | MICs (mg/L) |  |  |  |  |  |  |  |  | Fold change |
| --- | --- | --- | --- | --- | --- | --- | --- | --- | --- | --- | --- |
|  |  | COL alone | COL + 1/4 AS +EDTA |  |  |  | COL + 1/8 AS +EDTA |  |  |  |  |
|  |  |  | 1/4 | 1/8 | 1/16 | 1/32 | 1/4 | 1/8 | 1/16 | 1/23 |  |
|  |  |  | EDTA | EDTA | EDTA | EDTA | EDTA | EDTA | EDTA | EDTA |  |
| JS | — | 0.25 | 0.015 | 0.125 | 0.125 | 0.125 | 0.03 | 0.25 | 0.25 | 0.25 | **0-17** |
| S34 | — | 0.125 | 0.008 | 0.015 | 0.015 | 0.015 | 0.02 | 0.0625 | 0.125 | 0.125 | **0-16** |
| S16 | — | 4 | 0.015 | 0.03 | 0.0625 | 0.0625 | 0.0625 | 0.125 | 0.125 | 0.5 | **8-267** |
| S20 | — | 2 | 0.0625 | 0.125 | 0.125 | 0.125 | 0.25 | 0.5 | 0.5 | 0.5 | **4-32** |
| S13 | + | 2 | 0.015 | 0.125 | 0.25 | 0.25 | 0.0625 | 0.25 | 0.5 | 0.5 | **4-133** |
| S30 | + | 4 | 0.015 | 0.0625 | 0.0625 | 0.125 | 0.125 | 0.125 | 0.25 | 0.5 | **8-266.6** |
| E16 | + | 2 | 0.00003 | 0.002 | 0.25 | 0.5 | 0.004 | 0.015 | 0.25 | 1 | **2-66667** |
| M15 | — | >64 | 32 | 32 | >64 | >64 | 32 | 32 | >64 | >64 | **0-2** |
| P01 | — | >64 | >64 | >64 | >64 | >64 | >64 | >64 | >64 | >64 | **0** |

Note: COL: colistin; AS: artesunate

To verify the antibacterial enhancement activity of AS and EDTA, the growth curves of *Salmonella* JS, S16 (*mcr*-*1*^-^), and S30 (*mcr*-*1*^+^) for the combined treatments were generated within 24 h (Figure 1). Generally, the higher concentration of COL (2 mg/L), either alone or in combinations, was found more effective against these strains than that of the lower concentration (0.1 mg/L). Compared with the control groups, when these bacteria were grown in the presence of COL alone (0.1 mg/L) or different drug combinations, the antimicrobial activity were not significant after incubation for 24 h (Figure 1a, b, c). By contrast, higher concentration of COL (2mg/L), alone or in combinations, showed better antibacterial activity, and a gradual increase in antibacterial activity was observed: AEC > AC > EC > C (Figure 1d, e, f). It’s worth noting that the effect of different combinations against *mcr*-*1*^+^ strain S30 was weaker than that of *mcr*-*1*^-^ S16 and standard sensitive strain JS.

**Figure 1.**
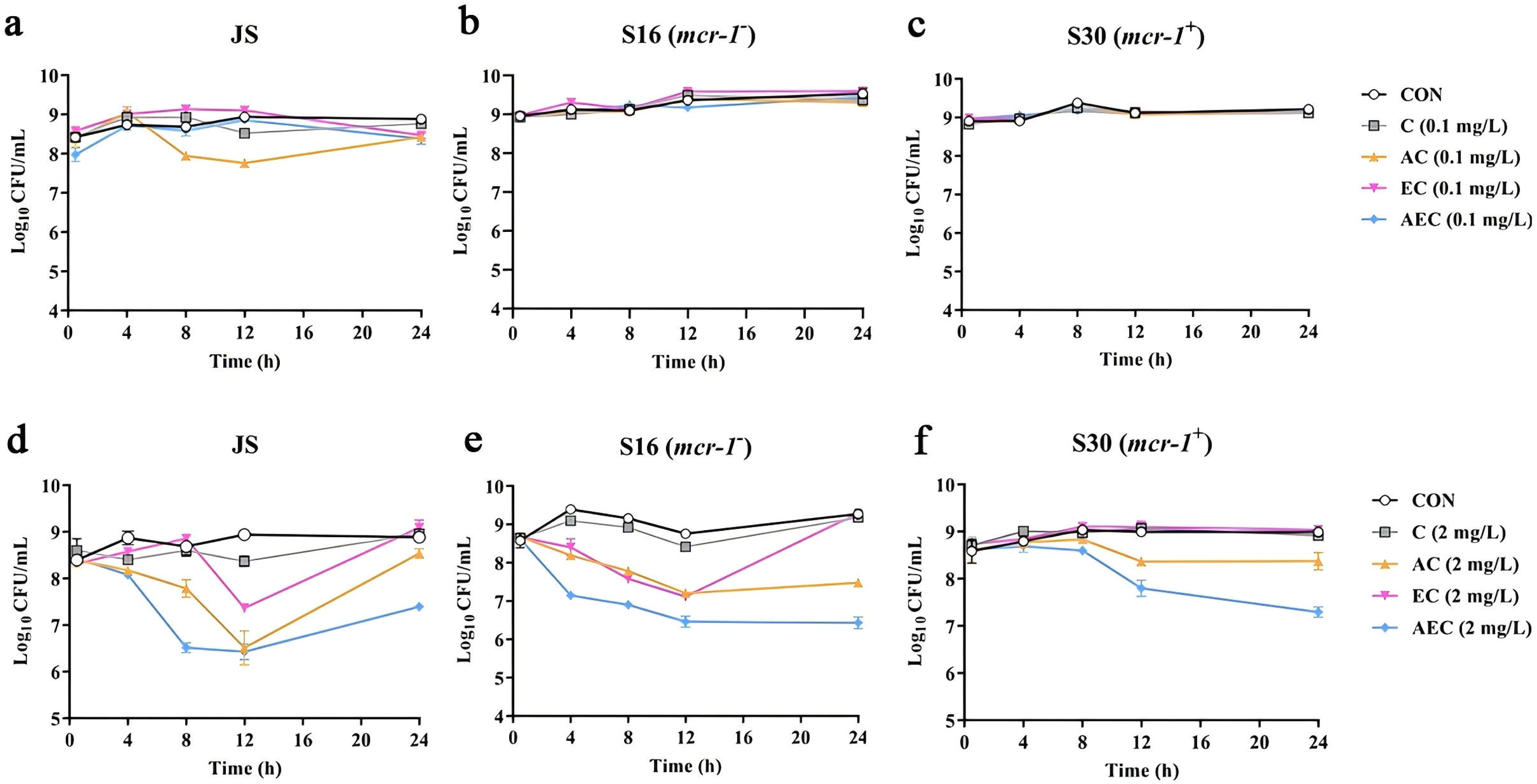
Time-kill curves of *Salmonella* strains JS, S16, and S30 with COL alone and in combinations. a-f Samples were treated with different concentration of COL (0.1 or 2 mg/L), alone or in drug combinations, for 12 h. When used in combination, 1/8 MIC of AS or EDTA was added to a final concentration of 156.3 or 15.6 mg/L, respectively. Counts of CFU/mL were performed on all cultures at each time point, and data are mean ± SD from representative of three independent experiments. CON indicates the negative control group.

### Artesunate and EDTA enhanced the membrane-damaging effect of colistin on *Salmonella*

In order to gain insight into the membrane-damaging bactericidal mechanism of COL alone or combinations, the damages to the bacterial outer and inner membrane (OM and IM) were severally monitored by measuring the fluorescent intensity of *Salmonella* strains S16 and S30 mixed with NPN and PI. Overall, the fluorescence signals of NPN and PI increased progressively with increases in the concentration of COL. After the treatment of AEC, S16 and S30 both showed strongest fluorescence signals of NPN, compared to those of other groups (Figure 2a, b). Nevertheless, AS treatment group also exerted a significant increase of fluorescent signal, although it is not as strong as that of AEC treated group. Meanwhile, there were rapid and significant increase in fluorescence of PI, when AEC or EC were added to bacterial cultures, and the two regimens played dominant roles in low or high concentrations of COL, respectively (Figure 2c, d). Therefore, these results indicated that the bacterial cell surfaces were severely damaged after the AEC treatment, via the rapid perturbations of OM and IM. Subsequently, the morphological changes of *Salmonella* strain S16 treated with different regimens were further investigated using Scanning electron microscope (SME) analysis to confirm the above membrane-damaging effect. As shown in Figure 3, SEM micro-graphs of control and solvent treated groups revealed that cells were short and rod-shaped, with rounded ends and intact cell membranes. As expected, after exposure to COL alone and different regimens, noticeable damage to the OM were observed, especially in AC and AEC treated group. Exposure to AC and AEC leading to cell damages characterized by folds, crevices, and depressions, which further suggested that AS and EDTA combined with COL resulted in remarkable bacterial membrane injury for antibacterial activity. Collectively, these data confirmed the membrane-damaging effects of AS, EDTA and COL combination against *Salmonella*.

**Figure 2.**
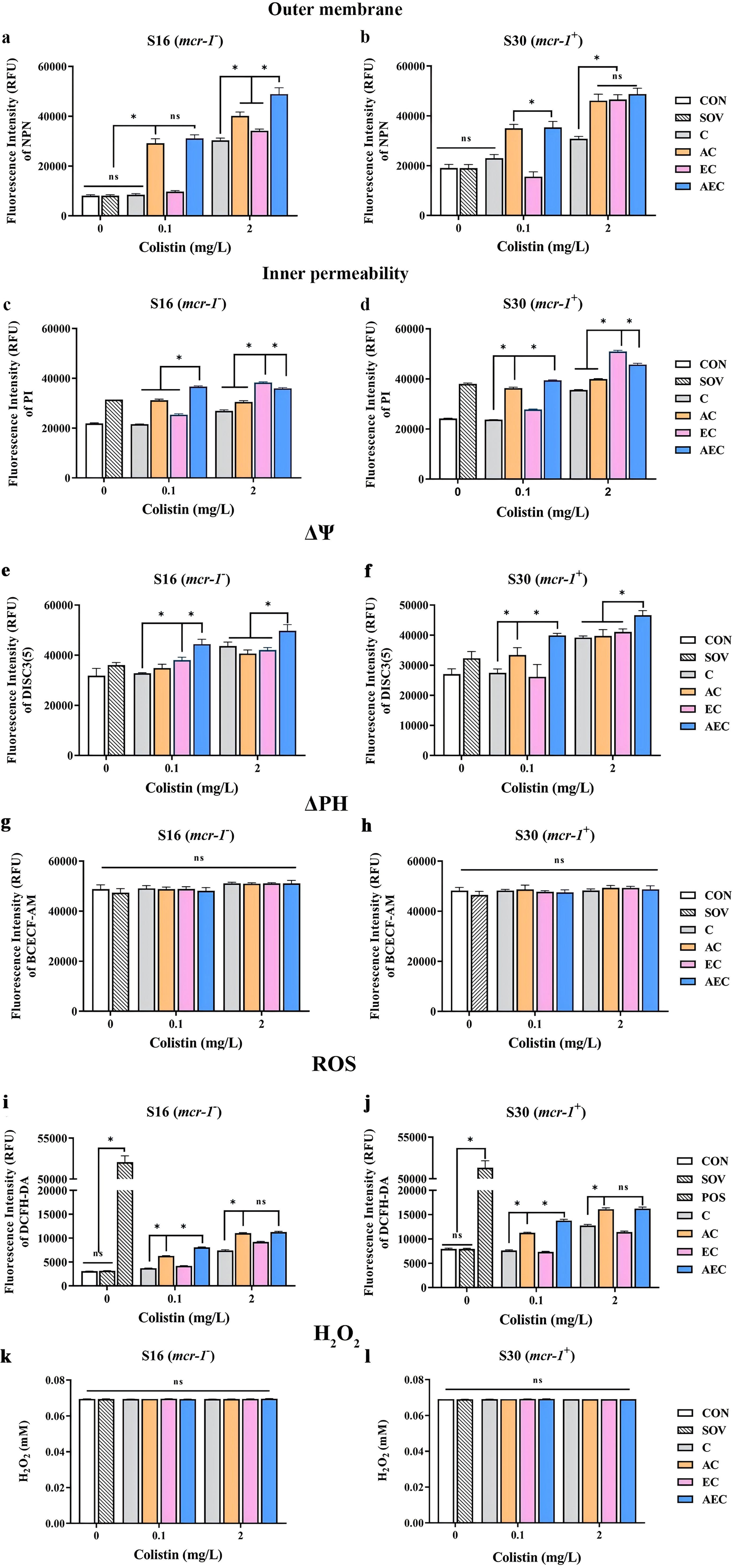
AS, EDTA, and COL affected membrane integrity, PMF, ROS, and H_2_O_2_ levels in S16 and S30 strains.. Different concentration of COL (0.1 or 2 mg/L) were used alone or in combination with AS and EDTA. When used in combination, 1/8 MIC of AS or EDTA was added to a final concentration of 156.3 or 15.6 mg/L, respectively. a-d AS, EDTA, and COL affected membrane integrity as measured by fluorescence probes NPN and PI. Error bars indicated standard deviations for 3 replicas (* *p* < 0.001, ns not significant). CON indicates the negative control group, and SOV indicates the solvent-exposed group. e-h Disruption of PMF is shown by measuring the dissipation of electric potential (Δψ) (a, b) and osmotic component ( Δ pH) (c, d). Error bars indicate standard deviations for 3 replicas (* *p* < 0.001, ns not significant). CON indicates the negative control group, and SOV indicates the solvent-exposed group. i-l Intracellular accumulation of ROS (i, j) and H_2_O_2_ (k, l) in S16 and S30 strains after 1 h treatment. Data were shown in the mean of triplicates ± SD (* *p* < 0.001, ns not significant). CON indicates the negative control group, SOV indicates the solvent-exposed group, and POS indicates the positive control group that treated with Rosup from the Total ROS Detection Kit.

**Figure 3.**
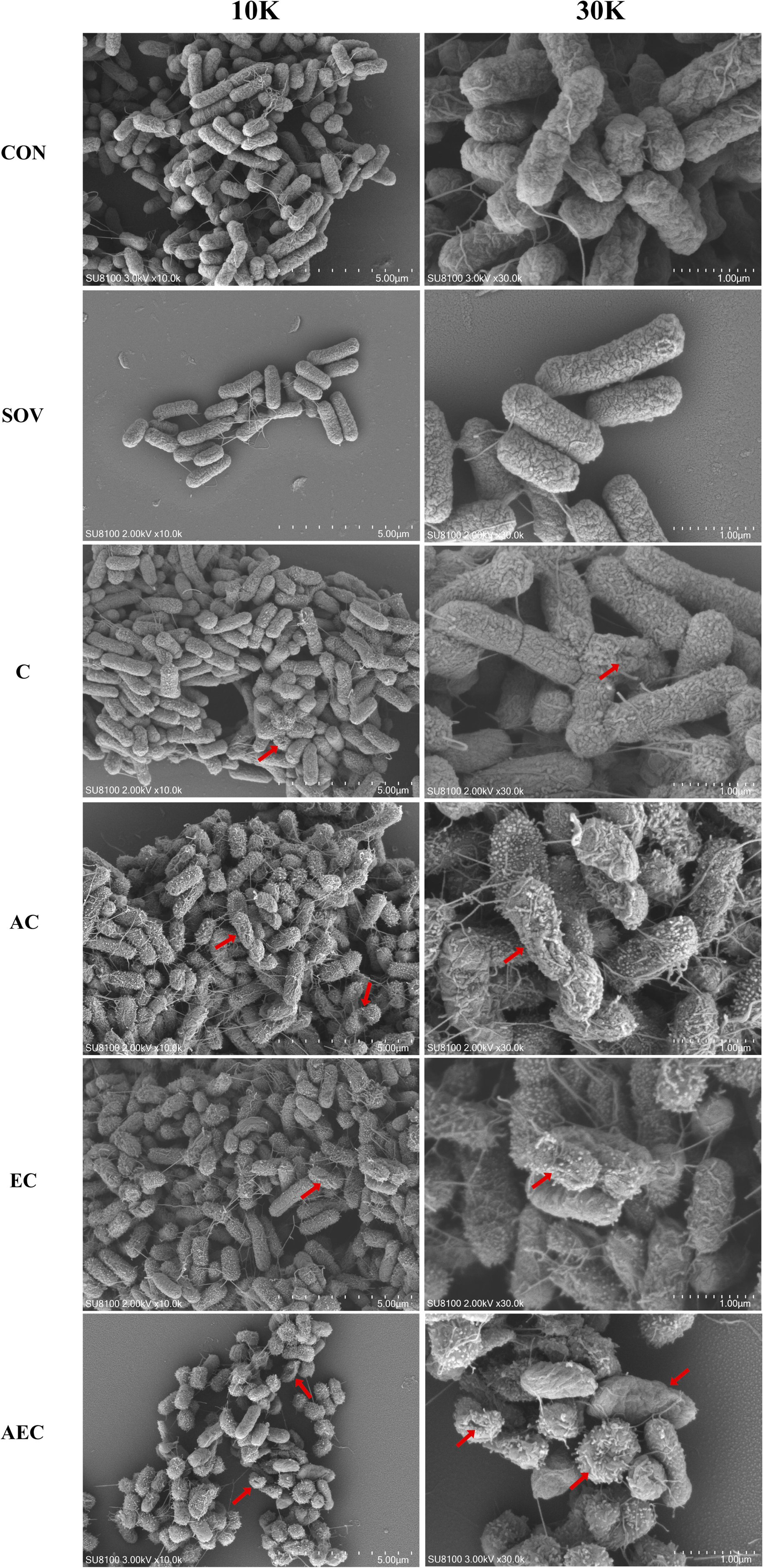
Morphological changes of S16 (*mcr*-*1*^-^) strain. The images were obtained after the treatment with COL (2 mg/L) alone or in combination with 1/8 MIC of AS (156.3 mg/L) or EDTA (15.6 mg/L). CON indicates the negative control group, and SOV indicates the solvent-exposed group. Red arrows indicate the cell damages characterized by folds, crevices, or depressions.

### AEC combination could collapse the Δψ component of Proton Motive Force (PMF) in *Salmonella*

In bacteria, the PMF, alternatively known as electrochemical proton gradient, is result from the extrusion of protons by the electron transport chain, and made up of the sum of two parameters: an electric potential (Δψ) and an osmotic component (ΔpH) (Farha *et al*, 2013). It has been reported to drives vital cellular processes in bacteria, including ATP synthesis, antibiotic transport, and cell division (Le *et al*, 2021). Therefore, PMF dissipation was considered as a promising strategy for combating microbial pathogens. In this work, we explored whether the antibacterial synergism activities of different combinations were accompanied by dissipation of PMF in cells. We uncovered that when compared to that of other groups, AEC treatment caused a rapid collapse of Δψ component, as shown by the increase in fluorescence values of DISC_3_(5) (Figure 2e, f). But none of these combinations observed significant dissipation of ΔPH, when compared to that of control group (Figure 2g, h). The above results indicated that AEC treatment was able to dissipate selectively the Δψ component of PMF.

### Other reactive oxygen species (ROS), not H_2_O_2_, contributed to the AS and EDTA-mediated efficacy enhancement of COL

ROS including superoxide (O^2-^), hydrogen peroxide (H_2_O_2_), and hydroxyl radical ( · OH), are commonly generated during the electron transfer process, which has been considered to associated with the lethal action of diverse antimicrobials. Subsequently, we investigated whether the addition of AS and EDTA could facilitate the intracellular ROS generation and stimulate the ROS-mediated killing. As shown in Figure 2i, j, compared to control group, total ROS increases in S16 and S30 strains were observed in AC and AEC groups either after 0.1 or 2 mg/L COL was added, but for COL alone and EC groups, the increase were only occured after the addition of a relatively high COL concentration (2 mg/L). In addition, the ROS accumulation level was further increased in AEC group, when extending the incubation time period to 6 h (Figure supplement 1). Moreover, we found that H_2_O_2_ did not contribute to the increase of total ROS, as there was no significant difference in intracellular level of H_2_O_2_ among all the groups Figure 2k, l. Collectively, these observations supported a role for ROS in AS and EDTA-mediated efficacy enhancement of COL.

### The transcriptome data exhibited more robust changes than that of metabolome among different comparison groups

A total of 6944 differentially expressed genes (DEGs) were performed KEGG pathway enrichment analysis, of which 1832 and 5112 transcripts were included in S16 and S30 strains, respectively (Figure supplement 2). Several canonical pathways, including two-component system (TCS), flagellar assembly and ABC transporters pathways, indicating similar directional changes in both strains were selected for further analysis, as shown in Figure 4. Since AEC incubation has displayed excellent antibacterial effects, we expected to screen the significantly differentially expressed genes (SDEGs) with similar variations in AEC vs. C, AEC vs. AC, and AEC vs. EC groups, and the SDEGs involved in TCS (*pagC*, *cheA*, *ompF*, *et al*.), flagellar assembly (*flgK*, *flgL*, *fliD*, *et al*) and ABC transporters (*oppuBB*, *osmX*, *gltI*, *dppA*, *et al*.) were selected (mostly down-regulated) and summarized in Figure 5.

**Figure 4.**
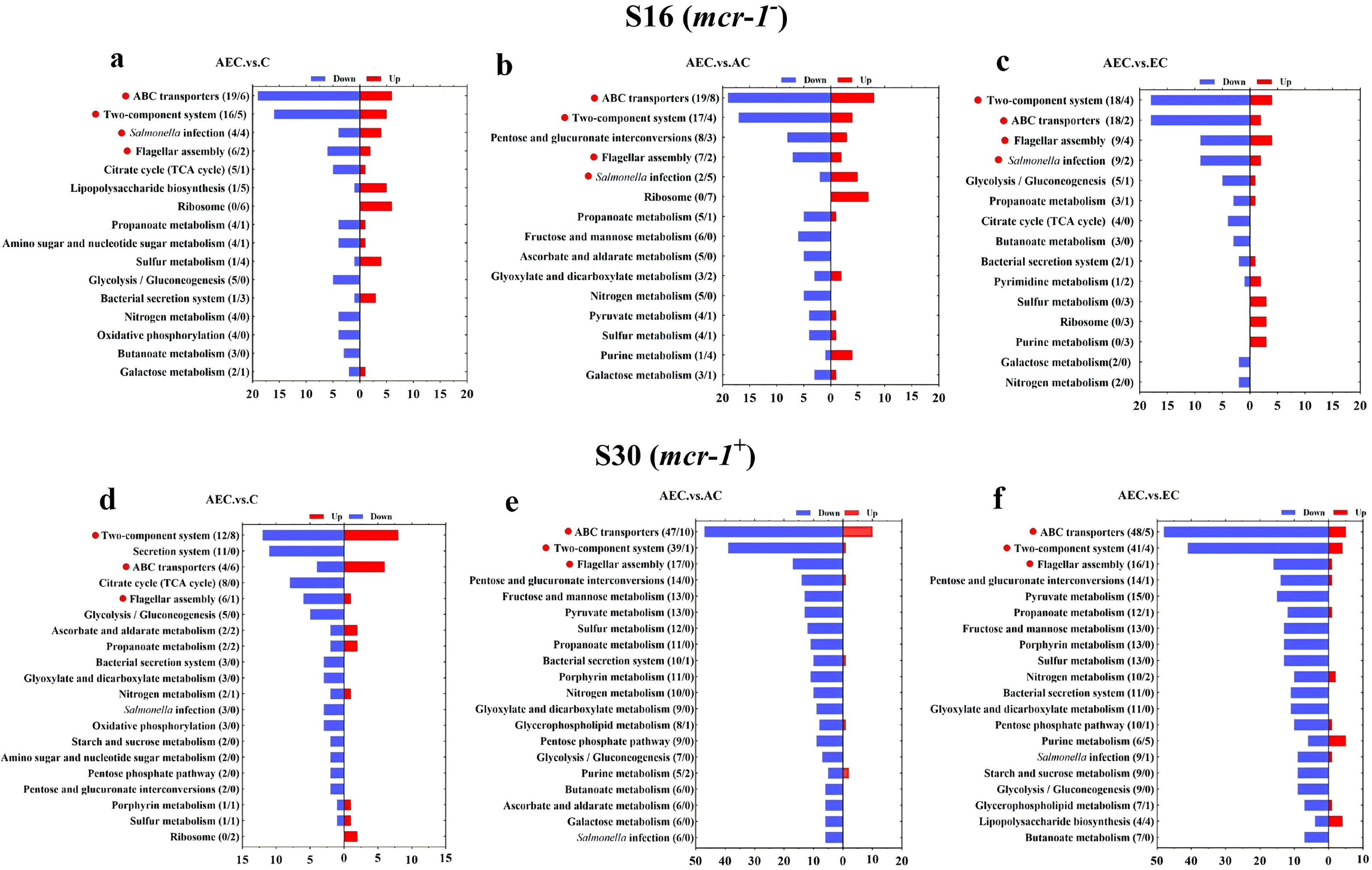
KEGG pathway analysis of SDEGs in S16 (a, b, c) and S30 (d, e, f) strains within the AEC .vs. C, AEC .vs. AC, and AEC .vs. EC groups. Samples were harvested after the trestment of COL (2 mg/L) alone or in combination with 1/8 MIC of AS (156.3 mg/L) or EDTA (15.6 mg/L) for 6 h. Pathway name and number of down-regulated (blue), up-regulated (red) genes in each pathway are indicated in parentheses on the left (down/up). Highlighted with red circles are the pathways that SDEGs mainly enriched and appeared simultaneously in different comparison groups.

**Figure 5.**
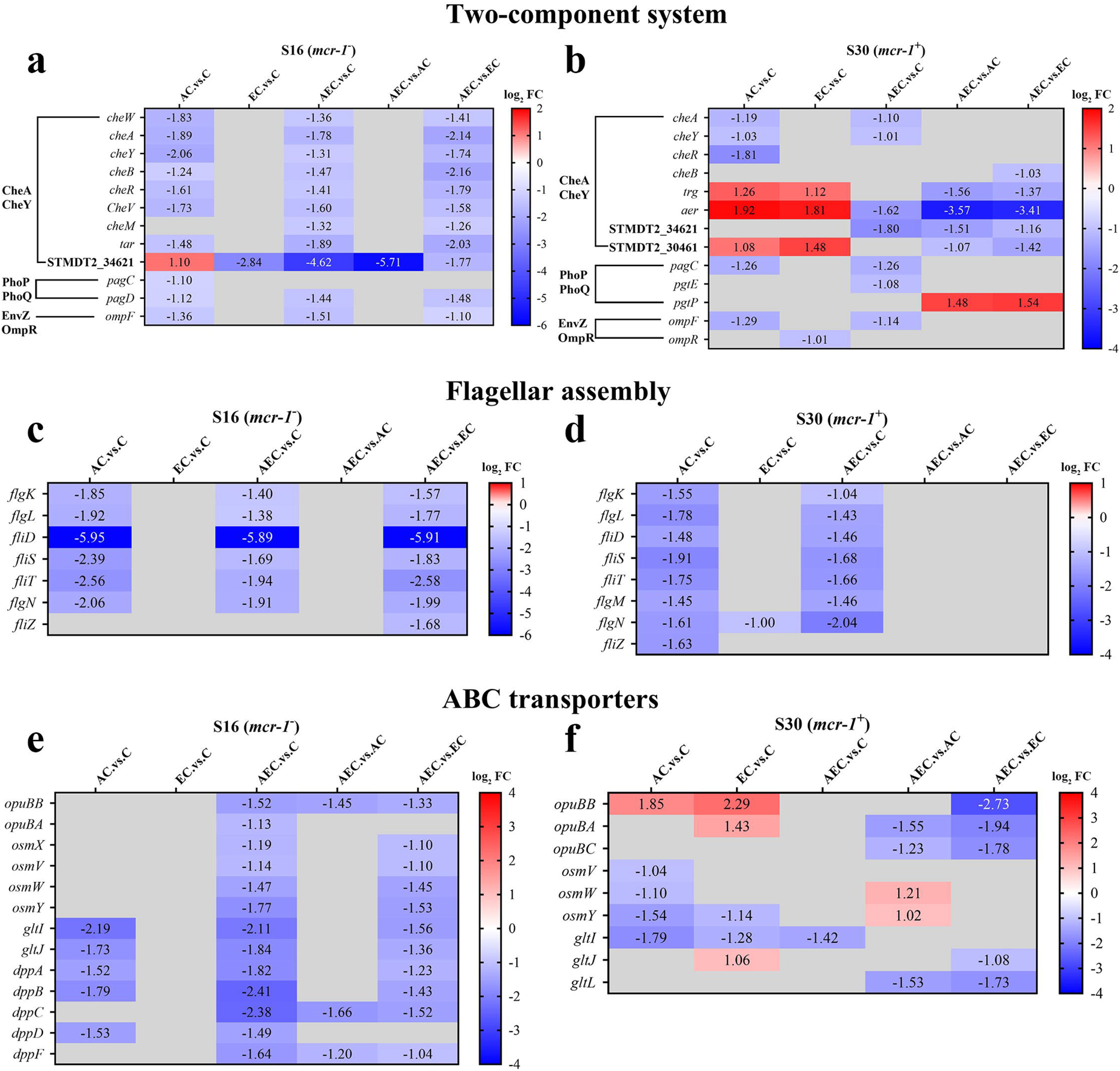
The SDEGs detected in two component system (a, b), flagellar assembly (c, d) and ABC transporters (e, f) pathways among different comparison groups, within S16 and S30 strains. Samples were harvested after the treatment of COL (2 mg/L) alone or in combination with 1/8 MIC of AS (156.3 mg/L) or EDTA (15.6 mg/L) for 6 h. Labels in each square indicate the log_2_ (fold change) of corresponding genes. Squares without label and gray background indicate the data are not credible (*p* > 0.05, |log_2_Fold Change| < 1.0). Background colors indicate the expression levels of the respective genes, red = up-regulated, blue = down-regulated. log_2_FC: log_2_Fold Change.

Unlike transcriptome changes, the metabolite alterations were much less abundant among AEC vs. C, AEC vs. AC, and AEC vs. EC groups, either in S16 or S30 strain. Unluckily, we demonstrated that there were low correlation between the metabolome and transcriptome data. According to the enrichment analysis, archidonic acid metabolism (down-regulated), degradation of aromatic compounds (up-regulated), taurine and hypotaurine metabolism (up-regulated) were the most prominent pathways showing differences primarily in AEC vs. C group (Figure 6).

**Figure 6.**
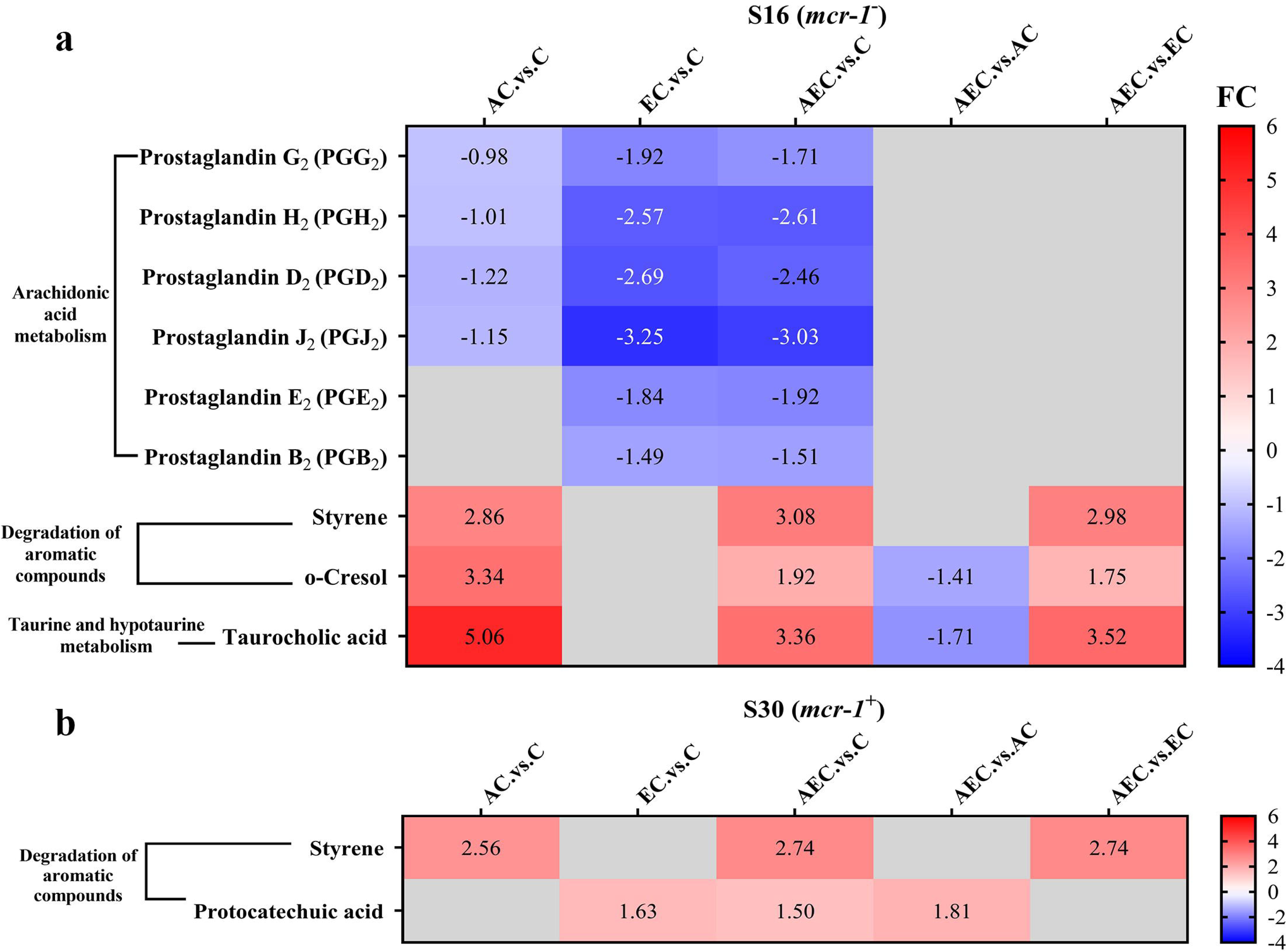
The SDMs detected in archidonic acid metabolism, degradation of aromatic compounds, taurine and hypotaurine metabolism pathways among different comparison groups, within S16 (a) and S30 (b) strains. Samples were harvested after the treatment of COL (2 mg/L) alone or in combination with 1/8 MIC of AS (156.3 mg/L) or EDTA (15.6 mg/L) for 6 h. Labels in each square indicate the fold changes of corresponding metabolites. Squares without label and gray background indicate the data are not credible (VIP < 1.0, 0.833 > Fold Change < 1.2 or Fold Change ≤ 0.833, *p* ≥ 0.05). Background colors indicate the fold changes of the respective metabolites, red = increased, blue = decreased.

### AS+EDTA+COL combination therapy is a promising therapeutic against *Salmonella* infection *in vivo*

The excellent bactericidal synergism against *Salmonella in vitro* of AEC combination further prompted us to confirm the effect *in vivo* for *Salmonella* S30 (*mcr*-*1*^+^) infected mouse models. Consistent with the synergistic bactericidal activity of AS, EDTA, and COL, the combination of AC (7.13 log_10_CFU/g Liver, 6.91 log_10_CFU/g spleen), EC (7.33 log_10_CFU/g Liver, 6.88 log_10_CFU/g spleen), and AEC (6.51 log_10_CFU/g Liver, 6.52 log_10_CFU/g spleen) outperformed single-drug treatments of COL (7.44 log_10_CFU/g Liver, 7.05 log_10_CFU/g spleen) and AS (7.54 log_10_CFU/g Liver, 7.02 log_10_CFU/g spleen). In particular, in the AEC combination treated samples, there were far fewer bacteria burden in spleen and liver when compared to other groups (Figure 7). These disparity *in vivo* further illustrate the synergy of AS and EDTA in combination with COL.

**Figure 7.**
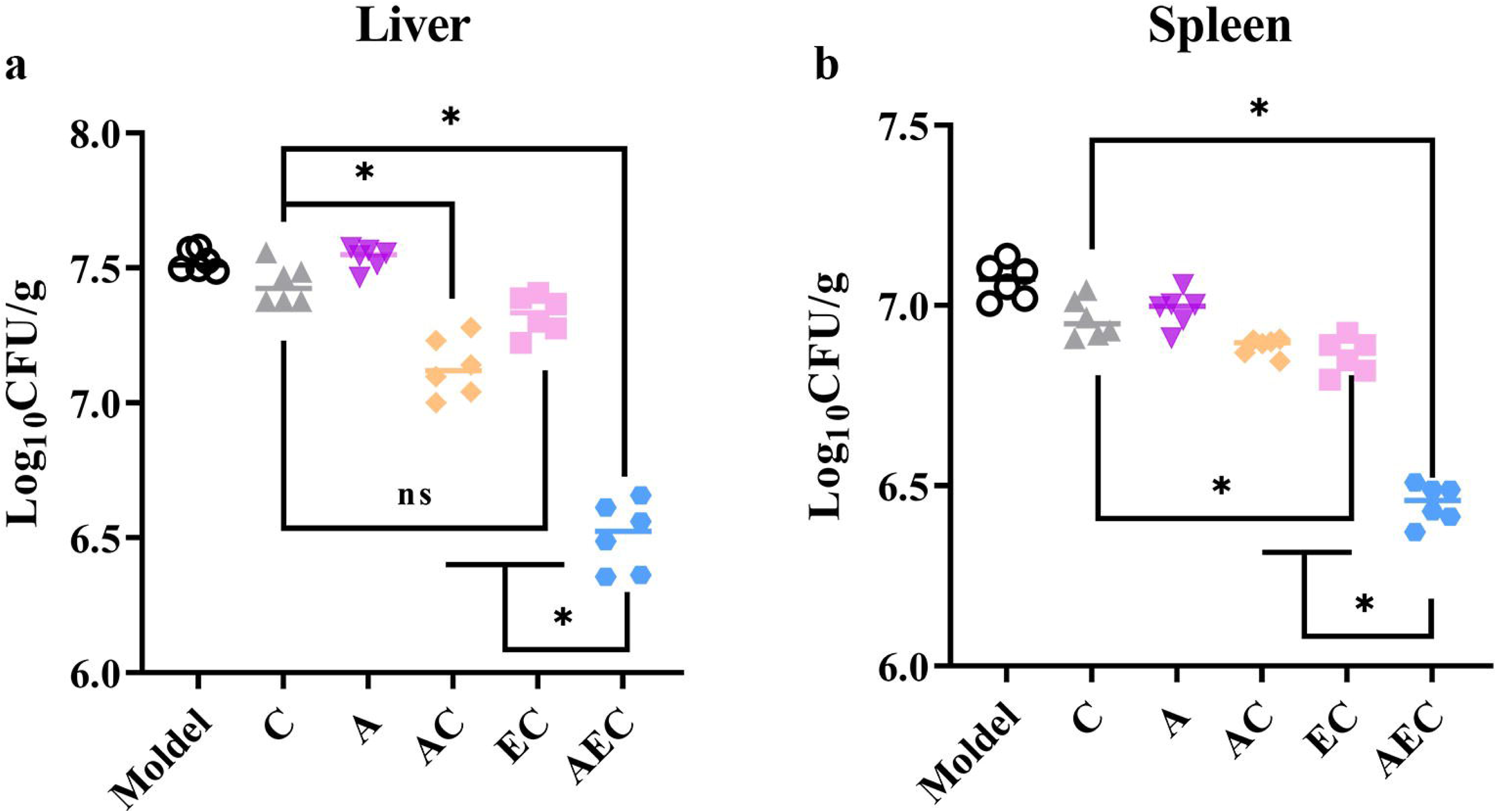
AS and EDTA potentiate colistin activity against *Salmonella* S30 (*mcr*-*1*^+^) *in vivo*. Kunming mice (n=6 per group) were intraperitoneally given a non-lethal dose of *Salmonella* S30 (1.31 × 10^5^ CFU), then treated with PBS, COL (10 mg/kg), AS (15 mg/kg), AS (15 mg/kg) + COL (10 mg/kg), EDTA (50 mg/kg) + COL (10 mg/kg), and AS (15 mg/kg) + EDTA (50 mg/kg)+ COL (10 mg/kg) by intraperitoneal injection. Bacterial loads were determined in spleen and liver and bacterial counts were computed and presented as the mean ± SD log_10_ CFU/mL. The *p* values were determined by one-way ANOVA (* *p* < 0.001, ns not significant).

## Discussion

### Membrane-damaging played a crucial role in the synergistic antimicrobial effects of AS+EDTA+COL combination

COL is an increasingly important antibiotic against serious infections caused by G^-^ bacteria. It damages both the OM and IM layers of the cell surface by targeting LPS, displacing cations that form bridges between LPS molecules, and thereby leading to disruption of the cell envelope and bacterial lysis (Sabnis *et al*, 2021). Our data showed that the AEC combination could permit ingress of NPN fluorophore into the OM, as well as the membrane impermeant dye PI into the IM, which fluoresces upon contact with DNA in the bacterial cytoplasm (Figure 2a-d). Thus, it indicated AEC combination has punched holes in both the OM and IM of whole bacterial cells. The considerably deformed of cell membranes observed by SEM further supported the above results that the membrane integrity, and permeability were damaged after AEC incubation. Considering that EDTA is used as a complexing agent, we tried to explore if it could assist COL to destroy the bacterial membrane and reduce their survival by chelating cations that stabilize LPS and the outer membrane. Nevertheless, we found minimal changes in MICs of different combinations to S16 and S30 strains, after different cations (Na^+^, K^+^, Ca^2+^, Mg^2+^, Mn^2+^, Zn^2+^) were supplemented (Table supplement 2). In contrast, LPS treatment resulted in dose-dependent changes in MICs of different combinations to S16 and S30 strains (Table supplement 3), which indicated that LPS on cell membrane was a crucial target for AEC to injure cell membrane and exert prominent antibacterial effects.

Whilst it is commonly believed that COL acts against G^-^ bacteria by cell membrane lysis, we hypothesised that alterations in bacterial membrane lipid composition may also be a possible Achilles’ heel to increase efficacy of COL after the AEC incubation in this paper. The metabolomic results of AEC vs. C group in this study showed a significant decrease of prostaglandins (PGs) in arachdonic acid metabolism pathway (Figure 6). Arachidonic acid is a highly abundant long chain polyunsaturated fatty acid in vertebrates, which have been proposed to have antibacterial roles. Exogenous Arachidonic acid has been reported to readily incorporated into the synthesis pathways of membrane phospholipids, and exert detrimental effects on membrane integrity by perturbing membrane ordering, altering membrane composition and increasing fluidity (Eijkelkamp *et al*, 2018; MacDermott-Opeskin *et al*, 2022). PGs are lipid compounds derived from arachidonic acid, which has been demonstrate to enhance biofilm development and fungal load in the murine vaginae of *Candida albicans* (Ells *et al*, 2011). Therefore, we speculated that the down-regulated lipid compounds PGs may lead to perturbtion of membrane phospholipids in cell membranes and reduce microbial viability.

Accumulation of toxic compounds may also be held responsible for the membrane damage. We noted that there were significant accumulation of styrene in both S16 and S30 strains (Figure 6). Styrene is naturally present as a minor metabolite, that can be synthesized at low levels by several microorganisms, like *Pencillium camemberti* and members of the *Styracaceae* family (Yeh *et al*, 2022). However, styrene itself is toxic to most cell types, and its hydrophobic molecules could readily partition into bacterial membrane, resulting in membrane disruption and cell death (Lian *et al*, 2016). Additionally, we found that the taurocholic acid and protocatechuic acid were up-regulated respectively in S16 and S30 strains (Figure 6). Taurocholic acid is usually a major component of the selective culture medium, for example MacConkey agar, for G^-^ bacteria, which has also been confirmed as a secondary metabolite of marine isolates and the soil bacterium *Streptococcus faecium*. Sannasiddappa *et al* have proved that taurocholic acid were able to inhibit growth of *Staphylococcus aureus* through increasing membrane permeability and disruption of the PMF (Sannasiddappa *et al*, 2017). Protocatechuic acid has been demonstrated to exert antimicrobial effects by disrupting the cell membranes, preventing bacterial adhesion and biofilm formation (Bernal-Mercado *et al*, 2018; Stojković *et al*, 2013). Consequently, these toxic compounds were anticipated to accelerate the destruction of cell membrane and thus enhance the antibacterial activity.

### The CheA of chemosensory system and virulence-related protein SpvD were critical for the bacteriostatic synergistic effect of AEC combination

Flagellar motility is intimately connected to chemotaxis, biofilm formation, colonisation and virulence of many bacterial pathogens. It is generally regulated by a chemotactic signaling system, which enables their movement toward favorable conditions and invade their hosts (Bolton, 2015). The chemotaxis proteins CheA, CheW, CheY, methyl-accepting chemotaxis proteins (MCPs) have already been identified as core components and present in all chemotaxis systems (Minamino *et al*, 2022). The *Salmonella* flagellum is composed of about 30 different proteins, such as FliD (the filament cap), FlgK and FlgL (the hook-filament junction) *et al* (Minamino *et al*, 2022). Tellingly, we have found an extensive down-regulation of chemotaxis and flagellar assembly related genes in S16 and S30 strains after AEC incubation (Figure 5a-d). Meanwhile, the swimming motility of the strains were decreased after AEC incubation (Figure supplement 5). The overexpression of *cheA* in S16 strain, but not *cheY*, *STMDT2-34621*, *aer*, *fliD*, *fliT*, could lead to noticeable increases of MICs by 4 to 32 fold, after EC or AEC incubation (Table supplement 4). These results suggested that the down regulation of the central component of chemosensory system CheA may affect both chemotactic motility and general structure of flagellum, thus attenuating *Salmonella* survival after AEC treatment.

In addition to the above findings, we also noted that there were relatively large number of SDEGs, most of them were down-regulated, enriched in the ABC transports pathway in S16 and S30 strains after AEC incubation (Figure 5e, f). ABC transporters are a class of transmembrane transporters, which mediate uptake of micronutrients, including saccharides, amino acids, metal ions *et al* and have also been shown to protect bacteria from hazardous compounds (Nguyen & Götz, 2016; Wang *et al*, 2023). The *opuBB* and *opuBA* encode components of the ABC-type proline/glycine betaine transport system, and their up-regulation were proved to promote accumulation of proline that acted as an osmoprotectant (Dupre *et al*, 2019). Similarly, the OsmU osmoprotectant systems, consisting of OsmV, OsmW, OsmY, and OsmX, were identified to enable bacterial survival at high-osmolarity through the accumulation of glycine betaine (Frossard *et al*, 2012). The *gltI* gene encodes glutamate/aspartate transport protein, and its deletion in *E. coli* were shown to result in attenuated survival under antibiotics, acid, and hyperosmotic stressors (Niu *et al*, 2023). Furthermore, the dipeptide permease operon (*dpp*), especially *dppA*, has been reported as an essential enzyme for survival of *Mycobacteria tuberculosis* under nutrient starvation conditions, and was associated with reduced bacterial burden in chronically infected mice in knockout studies (Fernando *et al*, 2022). Nevertheless, the overexpression of *opuBB*, *gltI*, *dppB* and *dppC* genes caused no detectable change in MICs for S16 strain after AEC incubation (Table supplement 4). Thus, these observations suggest that overexpression of these SDEGs in ABC transporters were not sufficient enough for causing changes in COL susceptibility, and additional factors may be required for the excellent bactericidal activity of AEC.

Besides that, we also noticed that the expression levels of *Salmonella* infection related genes *sseJ*, *spvD* and ribosomal protein genes *rpmJ*, *rpmE* were all strikingly changed in S16 strain (Figure supplement 4a). SseL and SpvD are effectors of *Salmonella* pathogenicity islands 1 and 2 (SPI1 and SPI2), which are required for full virulence during animal infections (Coombes *et al*, 2007; Grabe *et al*, 2016). The *rpmJ* gene codified a part of 50S ribosomal subunit, and its up-regulation has been related to size increase and slow growth of *E*. *coli* cells under starvation conditions (Peredo-Lovillo *et al*, 2019). The overexpression of *spvD* gene in S16 strain could attenuate the antibacterial activity of AC and AEC regimen with 4 fold increase in MICs (Table supplement 4), which indicated that the virulence-related protein SpvD contributed to the increase of COL susceptibility after AEC incubation.

### Artesunate can be considered as a potential MCR-1 inhibitor that enhances the efficacy of colistin

During our research process, several phenomenon have caught our attention, that the synergistic effect of AEC combination was irrelevant to whether the *mcr*-*1* gene exists or not, and the *mcr*-*1*^-^ strain S16 exhibited more robust changes than that of *mcr*-*1*^+^ strain S30 in different KEGG pathways. These results indicated that AS and EDTA were possible to exert synergistic effects by blocking the broad-spectrum resistance mechanisms (eg. efflux pumps, membrane damage), and coupling with the drug-specific resistance mechanisms (eg. MCR-1, β-lactamase). Previously, AS was capable of significantly enhancing the antibacterial activity of β-lactam antibiotics against *E*. *coli*, via inhibition of the efflux pumps such as AcrB, NorA, NorB, and NorC (Jiang *et al*, 2013; Li *et al*, 2011). Molecular docking experiments showed that AS could dock into AcrB very well forming five hydrogen bonds with Ser46, Gln89 and Gln176 (Wu *et al*, 2013). Whereas, in this paper, the inhibitory actions of different drug combinations on efflux pump were only observed in *mcr*-*1*^-^ strain S16, regardless of whether AS was added (Figure supplement 4c). Therefore, we hypothesised that AS may exert synergistic effects with COL against *mcr*-*1*^+^ S30 strain by targeting MCR-1 rather than efflux pump.

MCR-1 comprises two distinct domains, an N-terminal transmembrane domain and a soluble C-terminal α / β / α sandwich domain where the active site located. The active site contains a concentration of metal-binding residues to accommodate between one and four zinc ions (Wei *et al*, 2018). Several crystal structures of the soluble domain have been well studied, and six residues GLU246, THR285, HIS395, ASP465, HIS466, and HIS478 were found conserved among pEtN transferases (Ma *et al*, 2016). Especially the THR285 residue, which was highly conserved and providing a distinct electronegative potential to attract and bind the substrate pEtN (Son *et al*, 2019). Mutations of these residues and stripping the metals by EDTA could re-establish polymyxin B antibacterial action (Hinchliffe *et al*, 2017; Hu *et al*, 2016; Stojanoski *et al*, 2016). In this paper, we firstly analyzed the relative expression of *mcr*-*1* in S30 strain after the incubation of different drug combinations, and found a striking down-regulation of *mcr*-*1* gene whether after AC, EC incubation or AEC incubation, when compered to that of A and E treatment (Figure supplement 4b). Additionally, we performed the molecular docking between AS and MCR-1 to predict if there were possible interactions, and found that AS could bind to the residues arrounding the reported key residues within MCR-1 of *E*. *coli*, forming seven hydrogen bonds with THR283, SER284, TYR287, PRO481, and ASN482 residues (Figure supplement 4e). The interaction between AS and MCR-1 was further proved by competitive inhibitory assays, that the binding of AS with MCR-1 was blocked after the addition of polypeptide P_u_ (containing unmutated THR283, SER284, and TYR287 sites) and lead to significant increases of MICs by 8 fold after AC or AEC treatment (Table supplement 5). Nevertheless, we also noticed that the blocking effect could not be removed by peptide P_m_ (containing mutated THR283, SER284, and TYR287 sites) when EDTA exists. We supposed that EDTA may chelate zinc ions that required for MCR-1 activity. Thus, we suggested that AS could be developed as the a MCR-1 inhibitor, and coupled with the chelating agent EDTA may favor to additionally magnify its inhibitory effect.

### A mixed blessing: the excellent antibacterial activity and potential toxicity of AEC combination

In the preceding years, several studies have reported on the synergistic effects of COL with different candidates, such as antimicrobial agents, natural compounds and synthetically prepared molecules (Cui *et al*, 2024, Yi *et al*, 2022). Nevertheless, the potential toxicity of these combination therapy will be the prime concern affecting their clinical application, and a similar concern has also been raised for the AEC combination in this study. Although *in vitro* studies have determined that with increasing dose of AS and EDTA, the antibacterial synergistic activity was gradually enhanced, and meanwhie, may also resulting in more toxic side effects. Thus, In our study, the 1/8 MICs of AS and EDTA were selected to ensure excellent antibacterial activity whereas minimize the potential toxicity. The toxic side effects of AEC combination may be most probably caused by COL, which is well known to present several adverse toxic effects, and the dose-dependent nephrotoxicity is the most reported (Lu *et al*, 2016, Visentin *et al*, 2017). Conversely, EDTA and AS may be low toxicity. When used in combination with COL or AB569, EDTA has been proved to synergistically overcome the COL-resistant *Klebsiella pneumonieae* and MDR *Acinetobacter baumannii* with the concentration of 12000 mg/L and 73 mg/L, respectively (Bari *et al*, 2023, Bogue *et al*, 2021). These concentrations were reported non-toxic to primary adult human skin (dermal) fibroblasts and other normal body cells, which may also apply to the lower concentrations of EDTA (15.6 mg/L) used in AEC combination (Shein *et al*, 2021). Additionally, AS has been found to exert multiple pharmacological actions including anti-malaria, anti-tumor, anti-viral and anti-inflammatory effects, which has displayed a relatively safe toxicity profile with the LD_50_ values being 4223 mg/kg (Cheong *et al*, 2020). However, the potential toxic effects of AEC combination is still unknown and requires further investigations.

In summary, our results established that the combination of COL with AS and EDTA was a promising candidate for combating infections caused by MCR-negative and -positive COL-resistant *Salmonella*. The membrane-damaging effect, accumulation of toxic compounds, inhibition of MCR-1 were supposed to play synergistic roles in reversing COL resistance of *Salmonella* (Figure 8). The CheA of chemosensory system and virulence-related protein SpvD were critical for the bacteriostatic synergistic effects of AEC combination. Selectively targeting CheA, SpvD or MCR using the natural compound AS could be further investigated as an attractive strategy for treatment of *Salmonella* infection.

**Figure 8.**
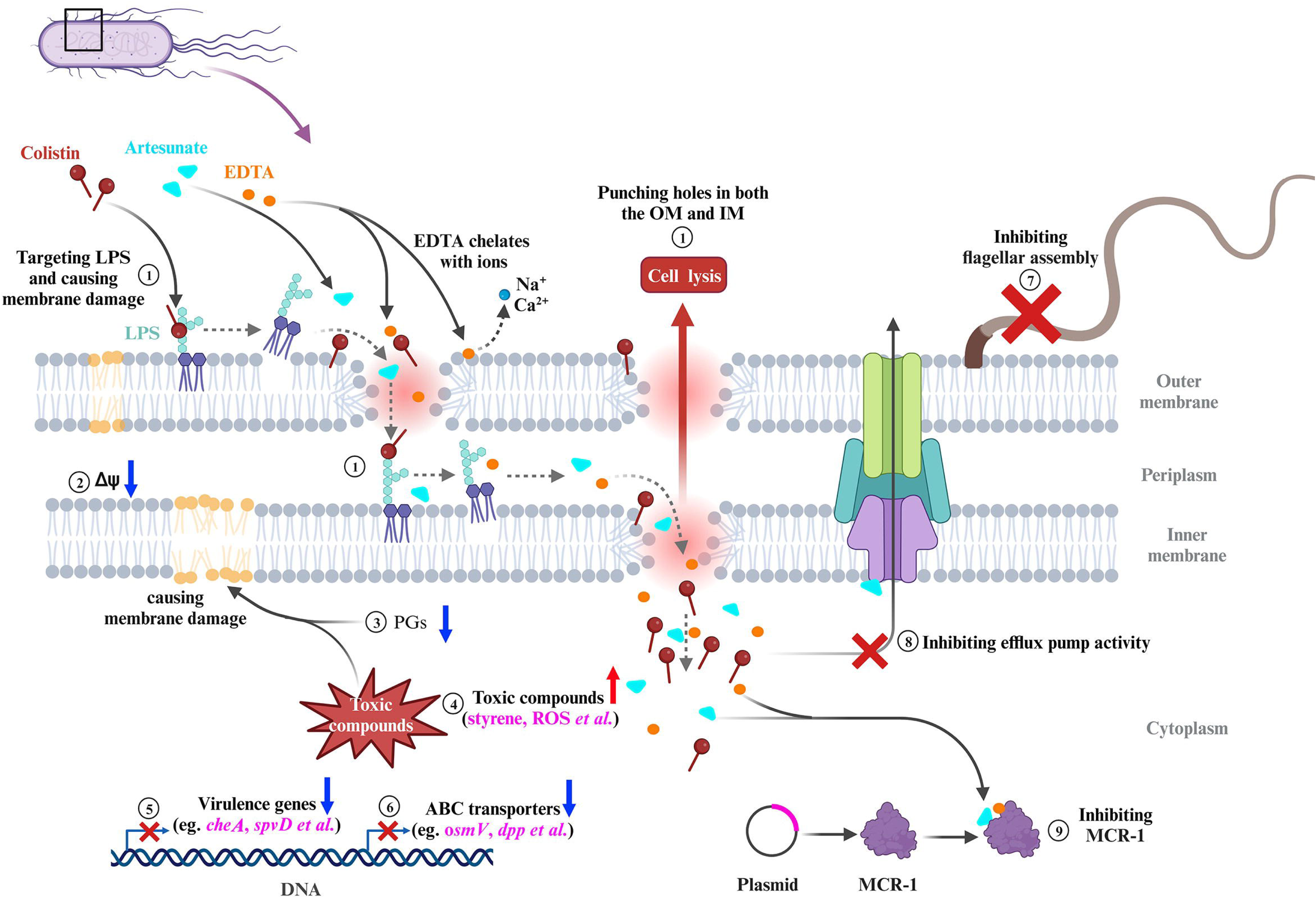
Scheme summarizing the proposed mechanisms that AS and EDTA enhancing the antibacterial effect of COL against *Salmonella*. ① COL and EDTA cause the membrane damages by targeting LPS and chelating cations, which punch holes in both the outer membrane (OM) and inner membrane (IM). ② AEC combination collapses the Δψ component of Proton Motive Force (PMF) in *Salmonella.* ③ The down-regulated lipid compounds prostaglandins (PGs) lead to perturbtion of membrane phospholipids in cell membranes. ④ Accumulation of toxic compounds (eg. styrene, ROS *et al*) could accelerate the destruction of cell membrane. ⑤ The down-regulation of chemotaxis, flagellar assembly, *Salmonella* infection related genes indicate impaired virulence of *Salmonella.* ⑥ The down-regulation of genes in ABC transporters indicate impaired stress tolerance of *Salmonella*. ⑦ AEC treatment results in the down-rugulation of flagellar assembly related genes and defective of flagellum. ⑧ AS was capable of significantly enhancing the antibacterial activities of antibiotics against *E*. *coli,* via inhibition of the efflux pumps. ⑨ AS could be developed as the a MCR-1 inhibitor, which may work synergistically with the EDTA chelation to inhibit MCR-1 and contribute to reverse the COL resistance of *mcr*-*1*-harboring *Salmonella* strains. FC: Fold Change.

## Materials and Methods

### Bacteria strains and agents

A total of 9 bacteria strains were used in this study (Table 1), including a multidrug-susceptible standard strain of *Salmonella* Typhimurium CVCC541 (named as JS), a COL-susceptible clinical strain of *Salmonella* (named as S34), four COL-resistant clinical strains of *Salmonella* (named as S16, S20, S13, and S30), a COL-resistant clinical strain of *Escherichia coli* (named as E16), and two strains of intrinsically COL-resistant species (*Morganella morganii* strain M15 and *Proteus mirabilis* strain P01). COL was purchased from Shengxue Dacheng Pharmaceutical, China. COL and EDTA-2Na were both dispersed into water at final concentration of 640 mg/L and 10000 mg/L, respectively. AS was purchased from Meilunbio (Dalian, China) and dissolved in water with 10% (V/V) N, N-dimethylformamide at a final concentration of 5000 mg/L. The 1-N-phenylnaphthylamine (NPN) was purchased from Sigma-Aldrich, USA. Propidium iodide (PI) was purchased from Thermo Fisher Scientific, USA. The 3,3-dipropylthiadicarbocyanine iodide DiSC_3_(5) and ethidium bromide (EtBr) were purchased from Aladdin, China. BCECF-AM, Reactive Oxygen Species Assay Kit, and Hydrogen Peroxide Assay Kit were purchased from Beyotime, China.

### Antibacterial activity *in vitro*

#### 1. Antimicrobial susceptibility testing

The MICs of COL, AS, and EDTA against all strains were determined by the 2-fold serial broth microdilution method according to CLSI guidelines (Wayne, 2021) The double and triple combination strategies were carried out as follow: AS (1/4, 1/8, or 1/16 MIC of AS)+ COL, EDTA (1/4, 1/8, or 1/16 MIC of EDTA) + COL, AS (1/4 MIC of AS)+ EDTA (1/4, 1/8, 1/16 or 1/32 MIC of EDTA) + COL, AS (1/8 MIC of AS)+ EDTA (1/4, 1/8, 1/16 or 1/32 MIC of EDTA) + COL. These medication strategies, including COL alone, AS + COL, EDTA + COL, AS + EDTA + COL, were abbreviated as C, AC, EC, and AEC, respectively. When determining the MICs of COL after drug combinations, AS or/and EDTA were pre-added into MHB broth with different final concentrations, namely 1/4, 1/8, 1/16 or 1/32 MIC of AS (312.5, 156.25, 78.13, 39.06 mg/L) or EDTA (for example 31.25, 15.63, 7.81, 3.91 mg/L for JS, S16 and S30 strains), then COL was added and diluted to make a 2-fold dilution series. The lowest concentrations with no visible growth of bacteria were defined as MIC values of COL after drug combinations.

#### 2. Time-kill assays

Time-kill assays were performed against the *Salmonella* strains JS, S16 (*mcr*-*1*^-^), and S30 (*mcr*-*1*^+^) with COL alone as well as in combinations (AC, EC, AEC). When combined with COL (0.1 or 2 mg/L), AS and EDTA were added at final concentrations equivalent to their 1/8 MICs. Overnight cultures were diluted 1:100 in fresh LB medium and grown to an OD_600_ of 0.5, then treated with different combinations for 24 h. The cultures were serially diluted 10-fold and spreaded over sterile nutrient agar at 0.5, 4, 8, 12 and 24 h. Bacterial colonies on individual plates were counted after overnight incubationat 37 ℃ and expressed as the log_10_ of colony forming units/mL (CFU/mL).

### Fluorescent probe-permeability assays

Overnight cultures of *Salmonella* strains S16 (*mcr*-*1*^-^) and S30 (*mcr*-*1*^+^) were diluted 1:100 in fresh LB medium and grown to an OD_600_ of 0.7. Cells were harvested and washed twice with PBS or HEPES, then re-suspended in the same buffer to OD_600_ ≈ 0.5 for further analysis. Different drug combinations or Fluorescent probes were added and incubated when necessary. The medication strategies used in the fluorescence probe assays were as follow: C, AC, EC, and AEC. The final concentration of COL was 0.1 or 2 mg/L, when used alone or in drug combinations. AS and EDTA were added at final concentrations equivalent to their 1/8 MICs, when used in drug combinations. Fluorescence intensity were measured with Spark 10M microplate spectrophotometer (Tecan, Switzerland).

#### 1. Cell membrane integrity assay

Bacterial suspensions in HEPES were mixed with either fluorescent probe 1-N-phenylnaphthylamine (NPN) or propidium iodide (PI) to a final probe concentrations of 10 μM for NPN or 15 μ M for PI. After incubation at 37 ℃ for 0.5 h, bacterial suspensions were then mixed with different drug combinations and incubated for another 1 h. Fluorescence measurements were then taken with the excitation wavelength at 350 nm (or 535 nm) and emission wavelength at 420 nm (or 615 nm) for NPN (or PI).

#### 2. Proton motive force assay

Bacterial suspensions in PBS were incubated with either 3,3-dipropylthiadicarbocyanine iodide (DiSC_3_(5), 0.5 μM) or pH-sensitive fluorescent probe BCECF-AM (20 μM) for 0.5 h to determine the membrane potential (Δψ) and pH gradient (ΔpH). Then these suspensions were mixed with different drug combinations and incubated for another 1 h. Finally, the fluorescence were measured with the excitation wavelength of 622 nm (or 488 nm) and emission wavelength of 670 nm (or 535 nm) for DiSC_3_(5) (or BCECF-AM).

#### 3. Total ROS measurement

The ROS-sensitive fluorescence indicator 2’, 7 ’-dichlorodihydro-fluorescein diacetate (DCFH-DA, 10 μM) was used to assess the ROS levels in bacterial cells. Bacterial suspensions in PBS were mixed with DCFH-DA and incubated for 0.5 h, then different drug combinations were added and incubated for another 1 h and 6 h. Finally, fluorescence intensity were measured at an excitation wavele of 488 nm and an emission wavelength of 525 nm.

#### 4. Efflux pump assay

Bacterial suspensions in PBS were incubated with ethidium bromide (EtBr) for 0.5 h, in a final concentration of 5 μM. Then different drug combinations were added and incubated for another 1 h, then the accumulation of EtBr in the cells were evaluated with excitation wavelength of 530 nm and barrier filter of 600 nm.

### H_2_O_2_ assay

The bacterial culture conditions and sample preparation were same as the fluorescent probe-permeability assays section. The luminescence absorbancy were measured by a Spapk 10 M Microplate reader (Tecan, Switzerland). The cellular H_2_O_2_ levels were assessed according to the kit procedure by using an Hydrogen Peroxide Assay Kit. Cells samples were prepared as described above and incubated with different drug combinations for 1.5 h. Cell precipitates were collected by centrifugation (10000 rpm), and 200 µL lysis solution were added under gentle shaking. Supernatants were then taken for luminescence (Abosorbance) detection at 560 nm.

### Scanning electron microscope (SME)

Overnight S16 (*mcr*-*1*^-^) culture was diluted 1:100 in fresh LB medium and grown to an OD_600_ of 0.7. Cells were then divided equally and different drug combinations were added, same as that in the fluorescent probe-permeability assays section. After 6 h incubation at 37 ℃, cells were washed three times with PBS and fixed with 2.5% glutaraldehyde at 4 ℃ for 24 h. Samples were then stained in 1% osmium tetroxide and dehydrated in a series of increasing ethanol concentrations (30% – 100%). The processed samples were dried in Critical Point Dryer (Quorum K850) and sputter-coated with gold. Finally, samples were observed and images were taken with SEM (HITACHI, SU8100).

### Omics analysis

Overnight cultures of S16 (*mcr*-*1*^-^) and S30 (*mcr*-*1*^+^) were diluted 1:100 in fresh LB medium and grown to an OD_600_ of 0.5. Cells were divided equally and different drug combinations were used, including C, AC, EC, and AEC. When combined with COL (2 mg/L), AS and EDTA were added at final concentrations equivalent to their 1/8 MICs. Samples were continually incubated 6 h, then washed three times with PBS and fast-frozen in liquid nitrogen for further use. Transcriptome and metabolome analysis were performed among different comparison groups, including AC vs. C, EC vs. C, AEC vs. AC, and AEC vs. EC, and carried out by Novogene Co. Ltd (Beijing, China).

#### 1. Transcriptome analysis

The clean reads were mapped to the *Salmonella* Typhimurium DT2 genome (HG326213.1) from NCBI using Bowtie2. The SDEGs were screened with a *p* value ≤ 0.05 and |log_2_Fold Change| ≥ 1. ClusterProfiler software were used to analysis the Gene Ontology functional enrichment (GO, https://geneontology.org/) or Kyoto Encyclopedia of Genes and Genomes pathway enrichment (KEGG, https://www.kegg.jp/kegg/pathway.html) of SDEGs.

#### 2. Metabolome analysis

In this study, Samples were analyzed by the non-targeted metabolomics with Liquid Chromatography-tandem Mass Spectrometry (LC-MS/MS) in either positive ion or negative ion mode. Compound Discoverer 3.1 software (Thermo Scientific, Waltham, MA, USA) was used for identification of metabolites based on the exact masses and fragmentation spectra. The significant differential metabolites (SDMs) were identified with VIP ≥ 1.0, Fold Change ≥ 1.2 or Fold Change ≤ 0.833, *p* value < 0.05. SDMs were then annotated and classified by KEGG Database (https://www.genome.jp/kegg/pathway.html), Human Metabolome Database (HMDB, https://hmdb.ca/metabolites), and Lipidmaps Database (https://www.lipidmaps.org/).

### Antibacterial activity *in vivo*

#### Bacterial preparation

Overnight culture of S30 (*mcr*-*1*^+^) was diluted 1:100 in fresh LB medium and grown to an OD_600_ of 0.7. Cells were harvested and washed twice with PBS, then adjusting the concentration of bacterial suspensions to 1.31 × 10^6^ CFU/mL for further use.

#### Animals and treatments

A total of 36 SPF Kunming mice (Six to eight-week-old, 18-22 g, half male and half female) were purchased from the Huaxing Experimental Animal Center of Zhengzhou (Zhengzhou, China) and divided into six groups (n = 6 per group): (1) PBS control group; (2) COL (10 mg/kg) group; (3) AS (15 mg/kg) group; (4) AS (15 mg/kg) + COL (10 mg/kg) group; (5) EDTA (50 mg/kg) + COL (10 mg/kg) group; (6) AS (15 mg/kg) + EDTA (50 mg/kg)+ COL (10 mg/kg) group. Each mice was intraperitoneally injected with 100 µL bacterial solution (1.31 × 10^5^ CFU). Treatment was initiated at 2 h post infection and continued for 3 days. Treatment was administered once per day by intraperitoneal injection according to the therapeutic dose mentioned above. Mices were all euthanized and spleen, liver of aseptic were collected, weighed, homogenized with PBS. Then the tissue homogenates were serially diluted with PBS in an appropriate amount and 100 μ L of each dilution was withdrawn and uniformly spread on SS agar plates. Bacterial counts were computed and presented as the mean ± SD log_10_ CFU/mL, after incubated at 37℃ for 16 to 18 h. Mice were maintained in a barrier facility and guaranteed strict compliance with the regulations for the Administration of Affairs Concerning Experimental Animals approved by the State Council of People’s Republic of China (11– 14–1988).The mouse experiments were approved by the Henan Science and Technology Department (protocol number SCXK 2019-0002).

### Data available

Transcriptome data have been submitted to the Sequence Read Archive database (SRA, https://www.ncbi.nlm.nih.gov/sra) under the BioProject accession number PRJNA1036120 (S16 strain) and PRJNA1036408 (S30 strain). Metabolome data have been submitted to the MetaboLights database (https://www.ebi.ac.uk/metabolights) under accession number MTBLS8875. These accession numbers have been released.

### Statistical analysis

Statistical analysis was conducted using Graphpad prism 9 and SPSS software. All data were derived from n ≥ 3 biological replicates and presented as mean ± SD. Without specific indication, differences between the independent groups (* *p* < 0.001) were assessed with student’s *t*-test or one-way ANOVA.

## Supporting information

Revised supplement materials

## Supplement materials

**Figure supplement 1.**
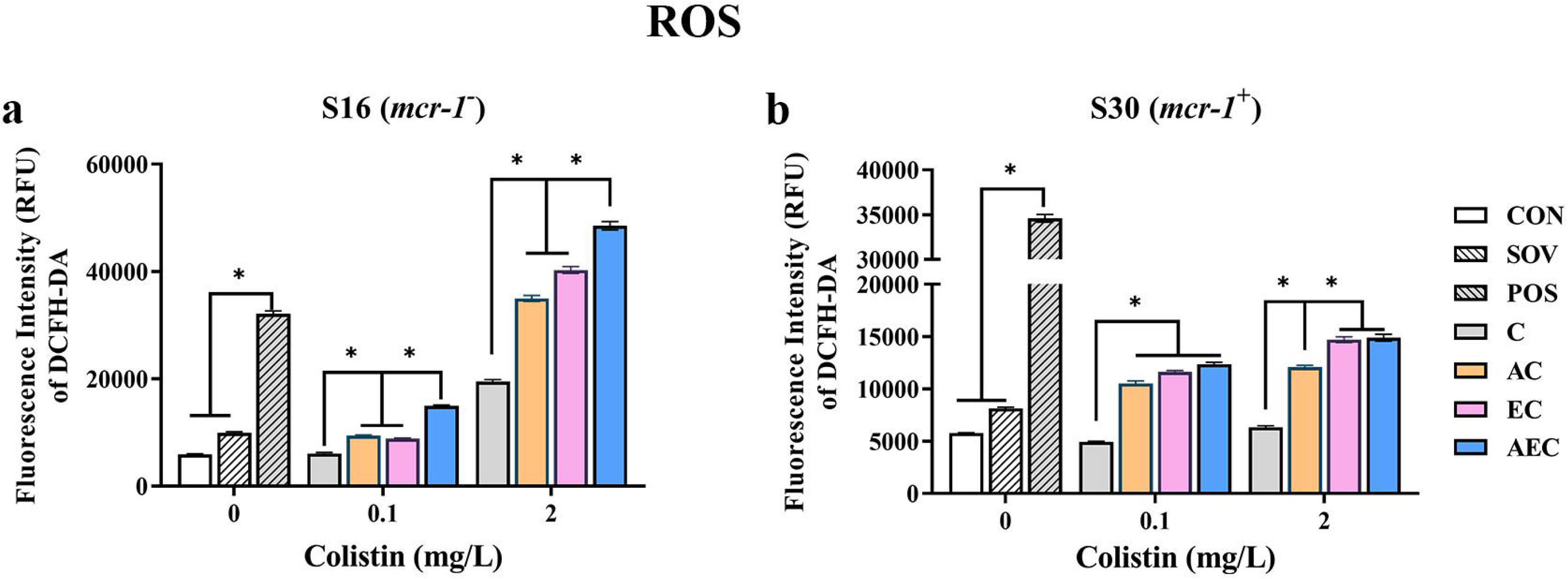
Intracellular accumulation of ROS in S16 and S30 strains after 6 h treatment.

**Figure supplement 2.**
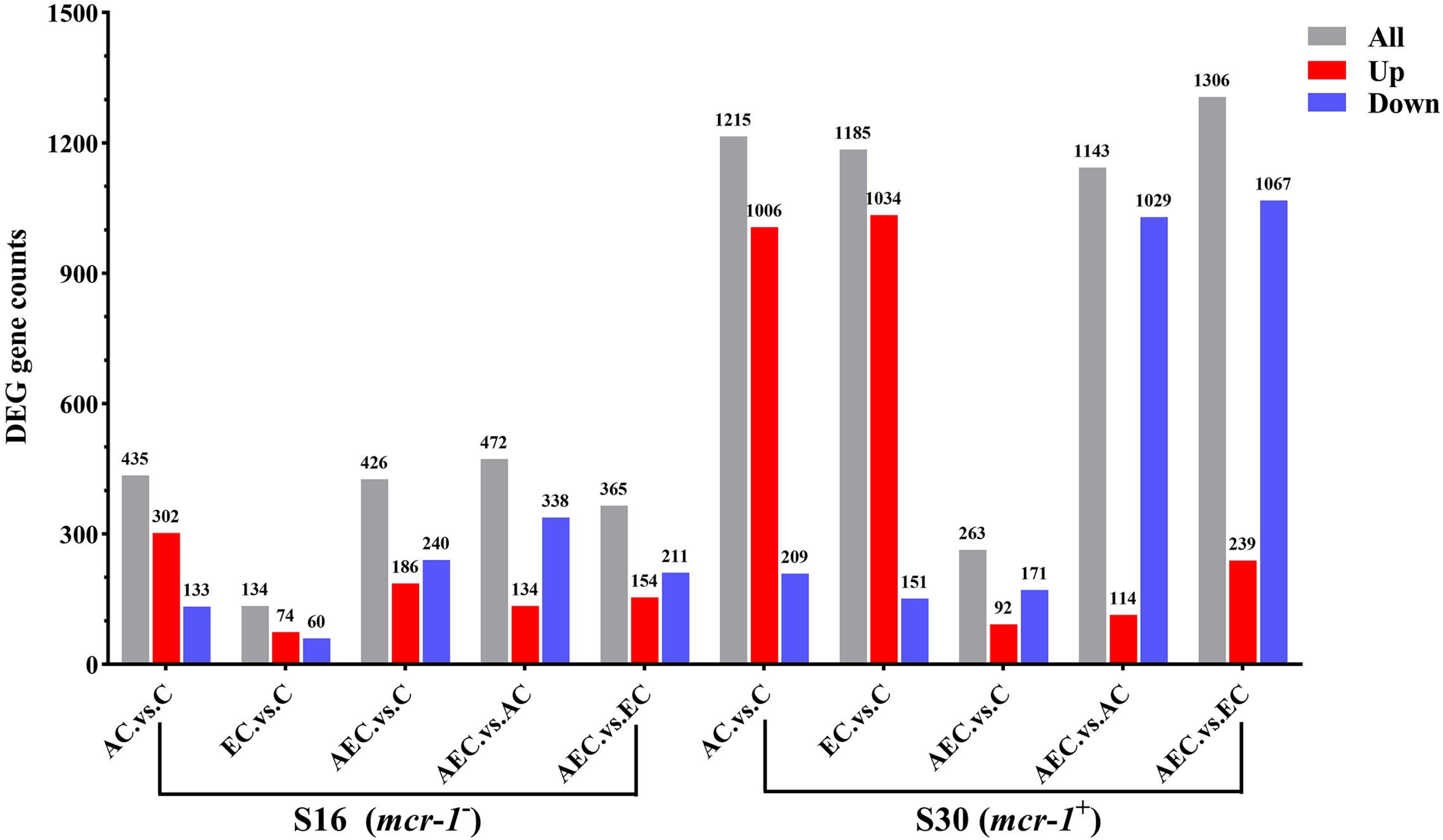
The number of DEGs are identified in S16 and S30 strains among different comparison groups.

**Figure supplement 3.**
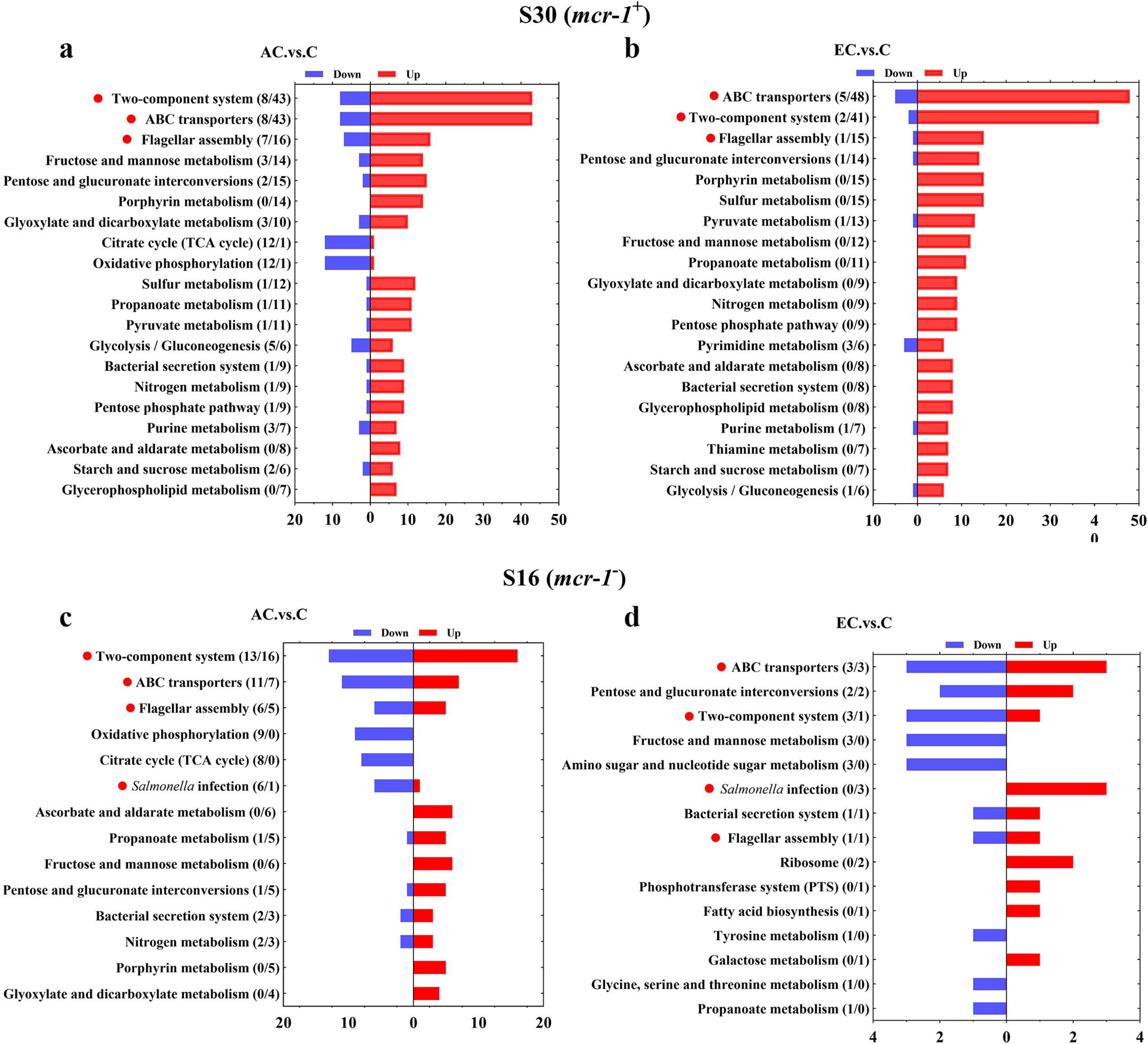
KEGG pathway analysis of SDEGs in S16 (a, b) and S30 (c, d) strains within the AC .vs. C, and EC .vs. C groups.

**Figure supplement 4.**
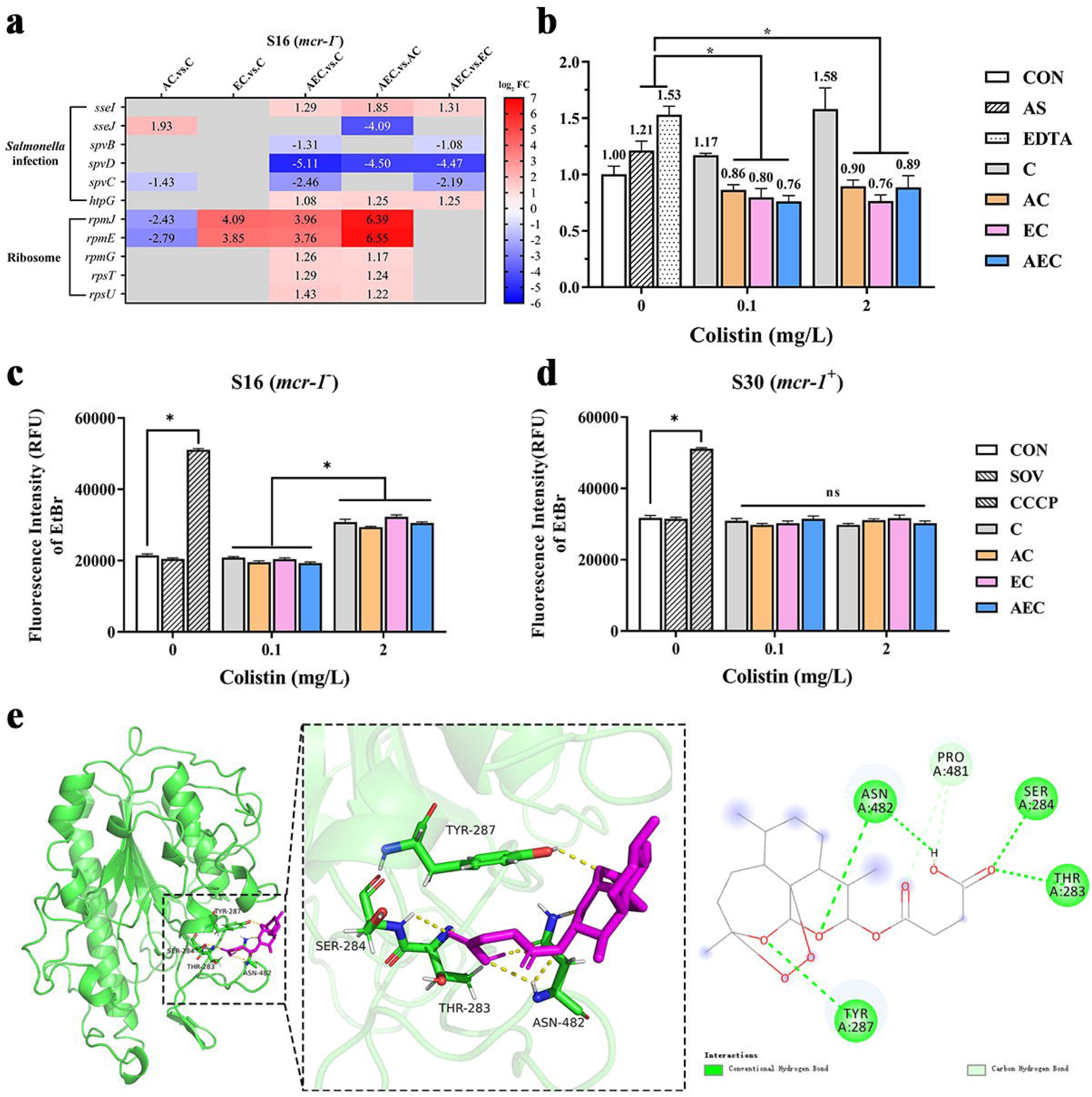
(a) The SDEGs detected in *Salmonella* infection and ribosome pathways; (b) Expression level of *mcr*-*1*; (c, d) efflux pump activity in S16 or S30 strain; (e) Putative pattern of interaction between AS and MCR-1 protein.

**Figure supplement 5.**
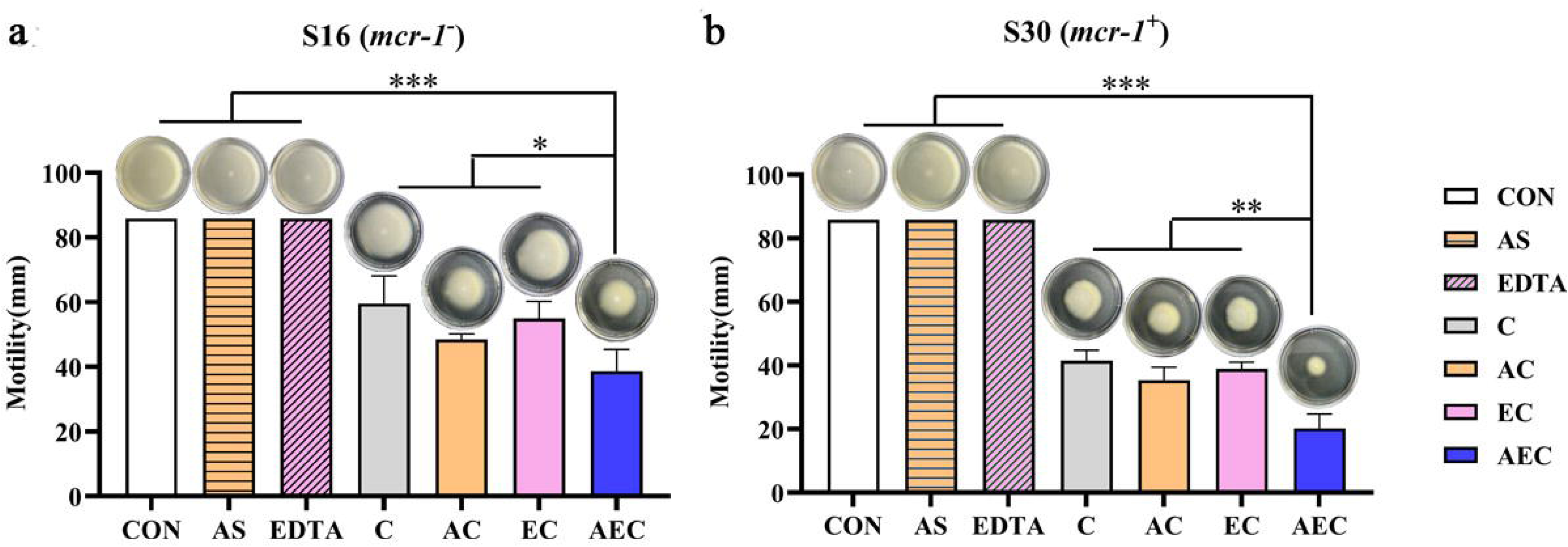
AS, EDTA, and COL inhibited the swimming motility of S16 (a) and S30 (b) strains.

**Table supplement 1.** The antibacterial activities of COL, AS and EDTA against the tested strains after single and double combinations

**Table supplement 2.** MIC values of COL against S16 and S30 strains after addition of exogenous cations

**Table supplement 3.** MIC values of COL against S16 and S30 strains after addition of exogenous LPS

**Table supplement 4.** MIC values of COL against S16 strain after the overexpression of different genes

**Table supplement 5.** MIC values of COL against S30 strain after the incubation of polypeptides

**Table supplement 6.** Sequences of primers used in this study

**Table supplement 7.** The antibacterial activities of COL, AS and EDTA against the tested strains after single and double combinations

**Table supplement 8.** The antibacterial activities of COL against the tested strains after single and triple combinations

**Table supplement 9.** The MICs of different antimicrobial drugs against the S16 and S30

**Supplement materials and methods**

## Acknowledgements

This work was supported by the National Natural Science Foundation of China (No. 32102716 and 32373069).

Ethical statement: This study was carried out in accordance with the guidelines of Henan Agricultural University Animal Ethics Committee.

## Author contributions

YZ and GH conceived the study. PL, XH, and CF performed experiments, results analysis and wrote the original draft. XC and QH analyzed the results and wrote the original draft. DH and XM analyzed the results. YZ, PL, and GH approved the final version of the manuscript.

## Data Availability

Data have been made available freely online.

## References

Aron Z and Opperman TJ (2016) Optimization of a novel series of pyranopyridine RND efflux pump inhibitors. Curr Opin Microbiol 33: 1–6.

Bari AK, Belalekar TS, Poojary A and Rohra S (2023) Combination drug strategies for biofilm eradication using synthetic and natural agents in KAPE pathogens. Front Cell Infect Microbiol 13: 1155699.

Bogue AL, Panmanee W, McDaniel CT, Mortensen JE, Kamau E, Actis LA, Johannigman JA, Schurr MJ, Satish L and Kotagiri N (2021) AB569, a non-toxic combination of acidified nitrite and EDTA, is effective at killing the notorious Iraq/Afghanistan combat wound pathogens, multi-drug resistant *Acinetobacter baumannii* and *Acinetobacter spp*. PLoS One 16: e0247513.

Bernal-Mercado AT, Vazquez-Armenta FJ, Tapia-Rodriguez MR, Islas-Osuna MA and Mata-Haro V (2018). Comparison of single and combined use of catechin, protocatechuic, and vanillic acids as antioxidant and antibacterial agents against uropathogenic *Escherichia Coli* at planktonic and biofilm levels. Molecules 23:2813.

Bolton DJ (2015). *Campylobacter* virulence and survival factors. Food Microbiol 48: 99–108.

Cui X, Liu X, Ma X, Li S, Zhang J, Han R, Yi K, Liu J, Pan Y, He D, Hu G, Zhai Y (2024) Restoring colistin sensitivity in colistin-resistant *Salmonella* and *Escherichia coli*: combinatorial use of berberine and EDTA with colistin. mSphere 9: e0018224.

Cheong DHJ, Tan DWS, Wong FWS and Tran T (2020) Anti-malarial drug, artemisinin and its derivatives for the treatment of respiratory diseases. Pharmacol Res 158: 104901.

Carroll LM, Gaballa A, Guldimann C, Sullivan G, Henderson LO and Wiedmann M (2019) Identification of novel mobilized colistin resistance gene *mcr*-*9* in a multidrug-resistant, colistin-susceptible *Salmonella enterica* Serotype Typhimurium isolate. mBio 10: e00853–19.

Coombes BK, Lowden MJ, Bishop JL, Wickham ME, Brown NF, Duong N, Osborne S, Gal-Mor O and Finlay BB (2007) SseL is a *Salmonella*-specific translocated effector integrated into the SsrB-controlled *Salmonella* pathogenicity island 2 type III secretion system. Infect Immun 75: 574–580.

Dupre JM, Johnson WL, Ulanov AV, Li Z, Wilkinson BJ and Gustafson JE (2019) Transcriptional profiling and metabolomic analysis of *Staphylococcus aureus* grown on autoclaved chicken breast. Food Microbiol 82: 46–52.

Eijkelkamp BA, Begg SL, Pederick VG, Trapetti C, Gregory MK, Whittall JJ, Paton JC and McDevitt CA (2018) Arachidonic acid stress impacts pneumococcal fatty acid homeostasis. Front Microbiol 9: 813

El-Sayed Ahmed MAE, Zhong LL and Shen C (2020) Colistin and its role in the Era of antibiotic resistance: an extended review (2000-2019). Emerg Microbes Infect 9: 868–885.

Ells R, Kock JL, Albertyn J, Kemp G and Pohl CH (2011) Effect of inhibitors of arachidonic acid metabolism on prostaglandin E₂ production by *Candida albicans* and *Candida dubliniensis* biofilms. Med Microbiol Immunol 200: 23–28.

Foletto VS, da Rosa TF, Serafin MB, Bottega A and Hörner R (2021) Repositioning of non-antibiotic drugs as an alternative to microbial resistance: a systematic review. Int J Antimicrob Agents 58: 106380.

Falagas ME and Kasiakou SK (2005) Colistin: the revival of polymyxins for the management of multidrug-resistant gram-negative bacterial infections. Clin Infect Dis 40: 1333–1341.

Farha MA, Verschoor CP, Bowdish D and Brown ED (2013) Collapsing the proton motive force to identify synergistic combinations against *Staphylococcus aureus*. Chem Biol 20: 1168–1178.

Fernando DM, Gee CT, Griffith EC, Meyer CJ, Wilt LA, Tangallapally R, Wallace MJ, Miller DJ and Lee RE (2022) Biophysical analysis of the *Mycobacteria tuberculosis* peptide binding protein DppA reveals a stringent peptide binding pocket. Tuberculosis (Edinb) 132: 102157.

Frossard SM, Khan AA, Warrick EC, Gately JM, Hanson AD, Oldham ML, Sanders DA and Csonka LN (2012) Identification of a third osmoprotectant transport system, the OsmU system, in *Salmonella enterica*. J Bacteriol 194: 3861–3871.

Galán-Relaño Á and Valero Díaz A (2023) *Salmonella* and Salmonellosis: An Update on Public Health Implications and Control Strategies. Animals (Basel) 13: 3666.

Gangathraprabhu B, Kannan S, Santhanam G, Suryadevara N and Maruthamuthu M (2020) A review on the origin of multidrug-resistant *Salmonella* and perspective of tailored *phoP* gene towards avirulence. Microb Pathog 147: 104352.

Grabe GJ, Zhang Y, Przydacz M, Rolhion N, Yang Y, Pruneda JN, Komander D, Holden DW and Hare SA (2016) The *Salmonella* effector SpvD is a cysteine hydrolase with a serovar-specific polymorphism influencing catalytic activity, suppression of immune responses, and bacterial virulence. J Biol Chem 291: 25853–25863.

Hinchliffe P, Yang QE, Portal E, Young T, Li H, Tooke CL, Carvalho MJ, Paterson NG, Brem J, Niumsup PR et al (2017) Insights into the mechanistic basis of plasmid-mediated colistin resistance from crystal structures of the catalytic domain of MCR-1. Sci Rep 7: 39392.

Hu M, Guo J, Cheng Q, Yang Z, Chan EWC, Chen S and Hao Q (2016) Crystal structure of *Escherichia coli* originated MCR-1, a phosphoethanolamine transferase for colistin resistance. Sci Rep 6: 38793.

Jiang W, Li B, Zheng X, Liu X, Pan X, Qing R, Cen Y, Zheng J and Zhou H (2013) Artesunate has its enhancement on antibacterial activity of β -lactams via increasing the antibiotic accumulation within methicillin-resistant *Staphylococcus aureus* (MRSA). J Antibiot (Tokyo) 66: 339–345.

Kaye KS, Pogue JM, Tran TB, Nation RL and Li J (2016) Agents of last resort: polymyxin resistance. Infect Dis Clin North Am 30: 391–414.

Laws M, Shaaban A and Rahman KM (2019) Antibiotic resistance breakers: current approaches and future directions. FEMS Microbiol Rev 43: 490–516.

Le D, Krasnopeeva E, Sinjab F, Pilizota T and Kim M (2021) Active efflux leads to heterogeneous dissipation of proton motive force by protonophores in bacteria. mBio 12: e0067621.

Li B, Yao Q, Pan XC, Wang N, Zhang R, Li J, Ding G, Liu X, Wu C, Ran D et al (2011) Artesunate enhances the antibacterial effect of β -lactam antibiotics against *Escherichia coli* by increasing antibiotic accumulation via inhibition of the multidrug efflux pump system AcrAB-TolC. J Antimicrob Chemother 66: 769–777.

Lian J, McKenna R, Rover MR, Nielsen DR, Wen Z and Jarboe LR (2016) Production of biorenewable styrene: utilization of biomass-derived sugars and insights into toxicity. J Ind Microbiol Biotechnol 43: 595–604.

Lu X, Chan T, Xu C, Zhu L, Zhou QT, Roberts KD, Chan HK, Li J and Zhou F (2016) Human oligopeptide transporter 2 (PEPT2) mediates cellular uptake of polymyxins. J Antimicrob Chemother 71: 403–412.

Lima WG, Alves MC, Cruz WS and Paiva MC (2018) Chromosomally encoded and plasmid-mediated polymyxins resistance in *Acinetobacter baumannii*: a huge public health threat. Eur J Clin Microbiol Infect Dis 37: 1009–1019.

Lin L, Nonejuie P, Munguia J, Hollands A, Olson J, Dam Q, Kumaraswamy M, Rivera H, Jr., Corriden R, Rohde M et al (2015) Azithromycin synergizes with cationic antimicrobial peptides to exert bactericidal and therapeutic activity against highly multidrug-resistant Gram-Negative bacterial pathogens. EBioMedicine 2: 690–698.

Ma G, Zhu Y, Yu Z, Ahmad A and Zhang H (2016) High resolution crystal structure of the catalytic domain of MCR-1. Sci Rep 6: 39540.

MacDermott-Opeskin HI, Panizza A, Eijkelkamp BA and O’Mara ML (2022) Dynamics of the *Acinetobacter baumannii* inner membrane under exogenous polyunsaturated fatty acid stress. Biochim Biophys Acta Biomembr 1864: 183908.

Minamino T, Kinoshita M and Namba K (2022) Insight into distinct functional roles of the flagellar ATPase complex for flagellar assembly in *Salmonella*. Front Microbiol 13: 864178.

Moya B, Barcelo IM, Bhagwat S, Patel M, Bou G, Papp-Wallace KM, Bonomo RA and Oliver A (2017) WCK 5107 (Zidebactam) and WCK 5153 are novel inhibitors of PBP2 showing potent “ β-Lactam Enhancer” activity against *Pseudomonas aeruginosa*, including multidrug-resistant metallo- β -lactamase-producing high-risk clones. Antimicrob Agents Chemother 61: e02529–16.

Nguyen MT and Götz F (2016) Lipoproteins of Gram-Positive bacteria: key players in the immune response and virulence. Microbiol Mol Biol Rev 80: 891–903.

Niu H, Li T, Du Y, Lv Z, Cao Q and Zhang Y (2023) Glutamate transporters GltS, GltP and GltI are involved in *Escherichia coli* tolerance in vitro and pathogenicity in mouse urinary tract infections. Microorganisms 11: 1173.

Pan X, Cen Y, Kuang M, Li B, Qin R and Zhou H (2020) Artesunate interrupts the self-transcriptional activation of MarA to inhibit RND family pumps of *Escherichia coli*. Int J Med Microbiol 310: 151465.

Peredo-Lovillo A, Dorantes-Alvarez L, Hernández-Sánchez H, Ribas-Aparicio R, Cauich-Sánchez P and Aparicio-Ozores G (2019) Functional properties and microstructure of *Leuconostoc citreum* and *Lactobacillus casei* shirota exposed to habanero pepper extract and inhibition of *Staphylococcus aureus*. Revista Mexicana de Ingeniería Química 18: 115–129.

Poirel L, Jayol A and Nordmann P (2017) Polymyxins: Antibacterial activity, susceptibility testing, and resistance mechanisms encoded by plasmids or chromosomes. Clin Microbiol Rev 30: 557–596.

Rosenthal PJ (2003) Antimalarial drug discovery: old and new approaches. J Exp Biol 206: 3735–3744

Sabnis A, Hagart KL, Klöckner A, Becce M, Evans LE and Furniss RCD (2021) Colistin kills bacteria by targeting lipopolysaccharide in the cytoplasmic membrane. eLife 10: e65836.

Sannasiddappa TH, Lund PA and Clarke SR (2017) *In vitro* antibacterial activity of unconjugated and conjugated bile salts on *Staphylococcus aureus*. Front Microbiol 8: 1581.

Son SJ, Huang R, Squire CJ and Leung IKH (2019) MCR-1: a promising target for structure-based design of inhibitors to tackle polymyxin resistance. Drug Discov Today 24: 206–216.

Stogios PJ, Spanogiannopoulos P, Evdokimova E, Egorova O, Shakya T, Todorovic N, Capretta A, Wright GD and Savchenko A (2013) Structure-guided optimization of protein kinase inhibitors reverses aminoglycoside antibiotic resistance. Biochem J 454: 191–200.

Stojanoski V, Sankaran B, Prasad BV, Poirel L, Nordmann P and Palzkill T (2016) Structure of the catalytic domain of the colistin resistance enzyme MCR-1. BMC biology 14: 81.

Stojković DS, Zivković J, Soković M, Glamočlija J, Ferreira IC, Janković T and Maksimović Z (2013) Antibacterial activity of *Veronica montana* L. extract and of protocatechuic acid incorporated in a food system. Food Chem Toxicol 55: 209–213.

Su F and Wang J (2018) Berberine inhibits the MexXY-OprM efflux pump to reverse imipenem resistance in a clinical carbapenem-resistant *Pseudomonas aeruginosa* isolate in a planktonic state. Exp Ther Med 15: 467–472.

Shein AMS, Wannigama DL, Higgins PG, Hurst C, Abe S, Hongsing P, Chantaravisoot N, Saethang T, Luk-In S, Liao T et al (2021). Novel colistin-EDTA combination for successful eradication of colistin-resistant *Klebsiella pneumoniae* catheter-related biofilm infections. Sci Rep 11: 21676.

Visentin M, Gai Z, Torozi A, Hiller C and Kullak-Ublick GA (2017) Colistin is substrate of the carnitine/organic cation transporter 2 (OCTN2, SLC22A5). Drug Metab Dispos 45: 1240–1244.

Wang J, Sheng Z, Liu Y, Chen X, Wang S and Yang H (2023) Combined proteomic and transcriptomic analysis of the antimicrobial mechanism of tannic acid against *Staphylococcus aureus*. Front Pharmacol 14: 1178177.

Wayne, PA (2021) Clinical and Laboratory Standards Institute. Performance standards for antimicrobial susceptibility testing: 31st ed. CLSI Supplement M100.

Wei P, Song G, Shi M, Zhou Y, Liu Y, Lei J, Chen P and Yin L (2018) Substrate analog interaction with MCR-1 offers insight into the rising threat of the plasmid-mediated transferable colistin resistance. Faseb j 32: 1085–1098.

Wei S, Yang Y, Tian W, Liu M, Yin S and Li J (2020) Synergistic activity of fluoroquinolones combining with artesunate against multidrug-resistant *Escherichia coli*. Microb Drug Resist 26: 81–88.

Wu C, Liu J, Pan X, Xian W, Li B, Peng W, Wang J, Yang D and Zhou H (2013) Design, synthesis and evaluation of the antibacterial enhancement activities of amino dihydroartemisinin derivatives. Molecules 18: 6866–6882.

Yeh V, Goode A, Johnson D, Cowieson N and Bonev BB (2022) The role of lipid chains as determinants of membrane stability in the presence of styrene. Langmuir 38: 1348–1359.

Yi K, Liu S, Liu P, Luo X, Zhao J, Yan F, Pan Y, Liu J, Zhai Y, Hu G (2022) Synergistic antibacterial activity of tetrandrine combined with colistin against MCR-mediated colistin-resistant *Salmonella*. Biomed Pharmacother 149: 112873.

Zhou Y, Liu B, Chu X, Su J, Xu L, Li L, Deng X and Li D (2022) Commercialized artemisinin derivatives combined with colistin protect against critical Gram-negative bacterial infection. Commun Biol 5: 931.

