## Supplementary material for "Artesunate, EDTA and colistin work synergistically against MCR-negative and -positive colistin-resistant *Salmonella*": Revised supplement materials

Running title: Synergy of artesunate, EDTA and colistin

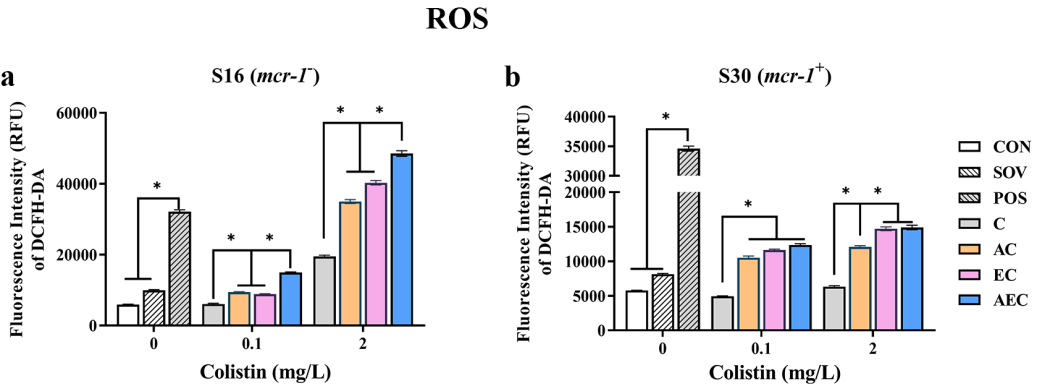

**Figure supplement 1. Intracellular accumulation of ROS in S16 and S30 strains after 6 h treatment.**

Different concentration of COL (0.1 or 2 mg/L) were used alone or in combination with 1/8 MIC of AS (156.3 mg/L) or EDTA (15.6 mg/L). Data were shown in the mean of triplicates ± SD (* *p* < 0.001). CON indicates the negative control group, and SOV indicates the solvent-exposed group.

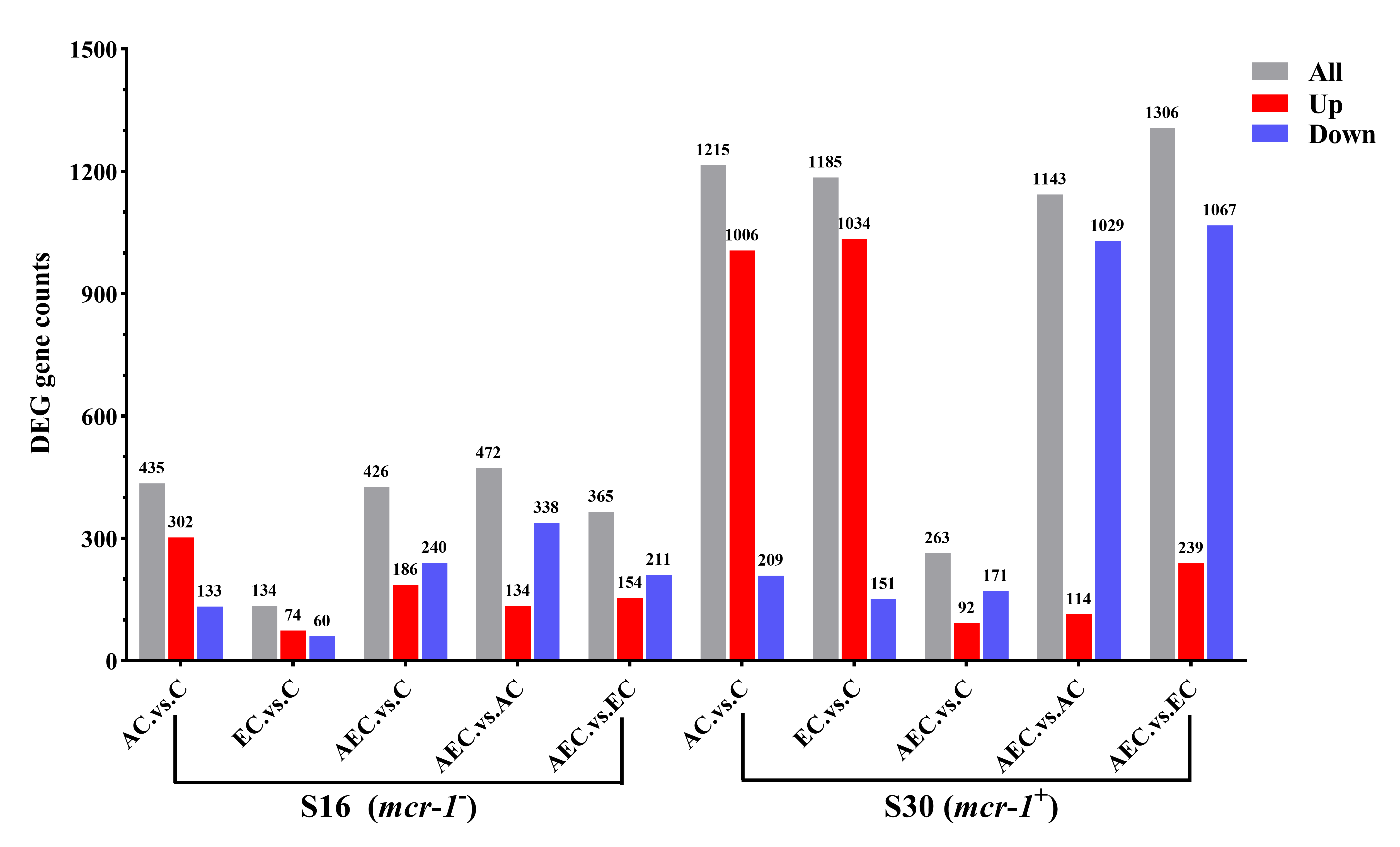

**Figure supplement 2. The number of DEGs are identified in S16 and S30 strains among different comparison groups.**

The total counts of DEGs are labeled at the top of each column and distinguished by different colors, all (gray), down-regulated (blue), up-regulated (red). Samples were harvested after the treatment of COL (2 mg/L) alone or in combination with 1/8 MIC of AS (156.3 mg/L) or EDTA (15.6 mg/L) for 6 h.

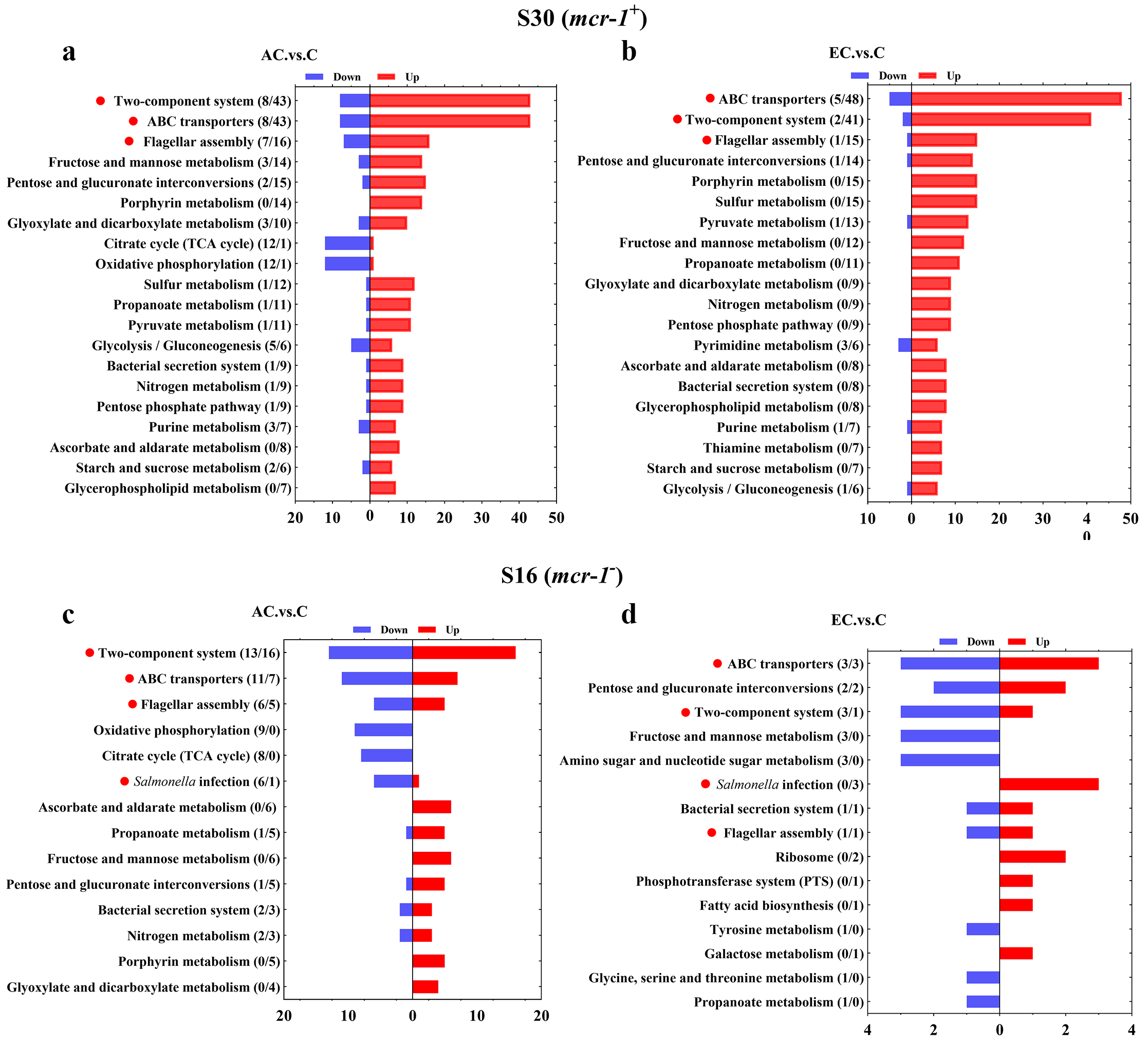

**Figure supplement 3. KEGG pathway analysis of SDEGs in S16 (a, b) and S30 (c, d) strains within the AC .vs. C, and EC .vs. C groups.**

a-d Samples were harvested after the treatment of COL (2 mg/L) alone or in combination with 1/8 MIC of AS (156.3 mg/L) or EDTA (15.6 mg/L) for 6 h. Pathway name and number of down-regulated (blue), up-regulated (red) genes in each pathway are indicated in parentheses on the left (down/up). Highlighted with red circles are the pathways that SDEGs mainly enriched and appeared simultaneously in different comparison groups.

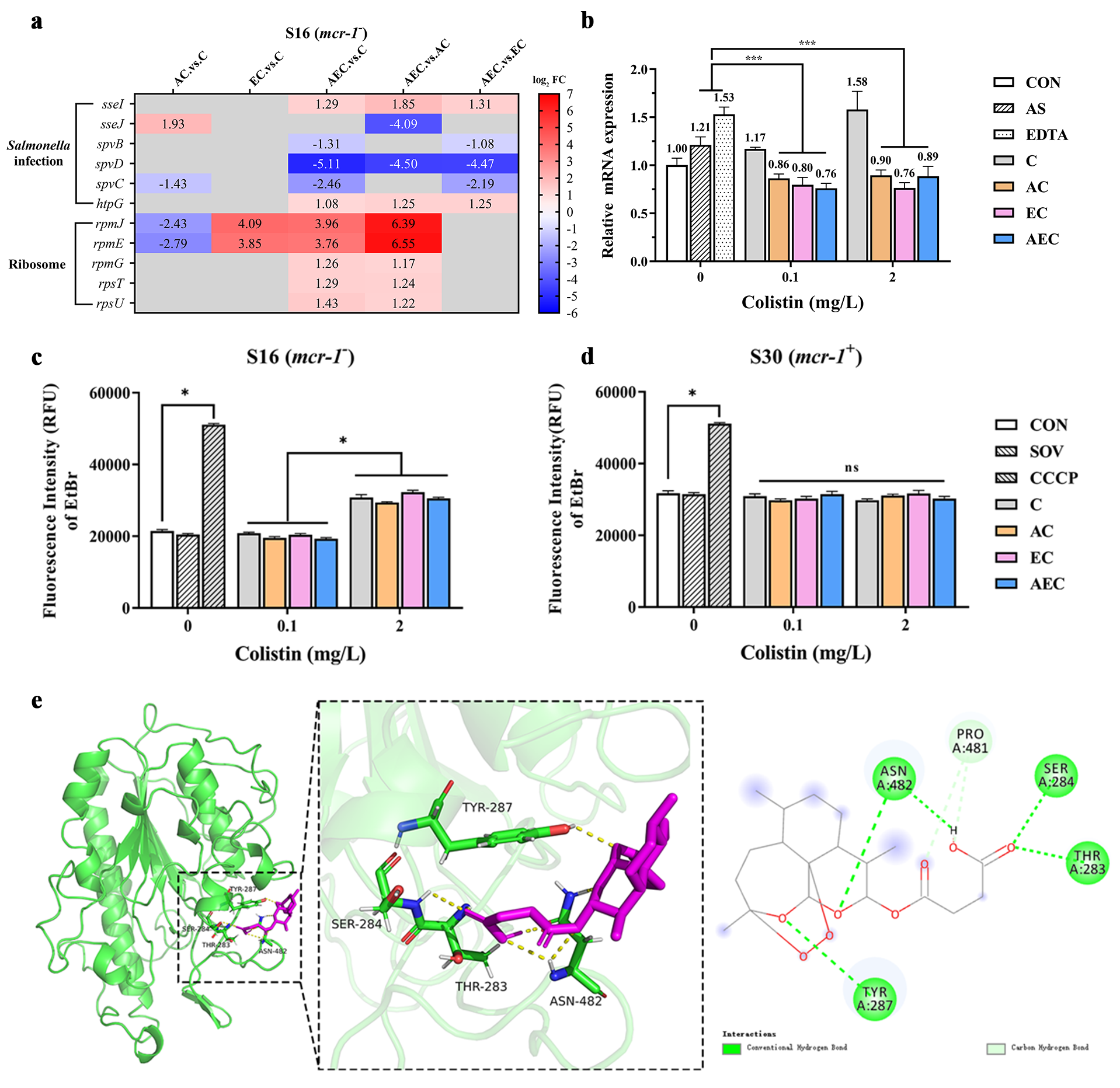

**Figure supplement 4.** **(a) The SDEGs detected in *Salmonella* infection and ribosome pathways; (b) Expression level of *mcr*-*1*; (c, d) efflux pump activity in S16 or S30 strain; (e) Putative pattern of interaction between AS and MCR-1 protein.**

Different concentration of COL (0.1 or 2 mg/L) were used alone or in combination with 1/8 MIC of AS (156.3 mg/L) or EDTA (15.6 mg/L).

a The SDEGs detected in *Salmonella* infection, ribosome pathways among different comparison groups, within S16 strain. Labels in each square indicate the log_2_ (fold change) of corresponding genes. Squares without label and gray background indicate the data are not credible (*p* > 0.05, |log_2_FoldChange| < 1.0). Background colors indicate the expression levels of the respective genes, red = up-regulated, blue = down-regulated. log_2_FC: log_2_Fold Change.

b	The relative expression of *mcr*-*1* in S30 strain after the incubation of different medication regimens. Significant differences were evaluated by student’s *t*-test analysis and shown with * *p* < 0.001.

c, d Intracellular accumulation of EtBr in S16 and S30 strains. Data were shown in the mean of triplicates ± SD (* *p* < 0.001, ns not significant). CON indicates the negative control group, and SOV indicates the solvent-exposed group.

e Putative pattern of interaction between AS and MCR-1 protein.The interactions formed between the amino acid residues (stick, green) and the docked AS molecule (stick, purple) in the MCR-1 binding sites are displayed in 3D and 2D views.

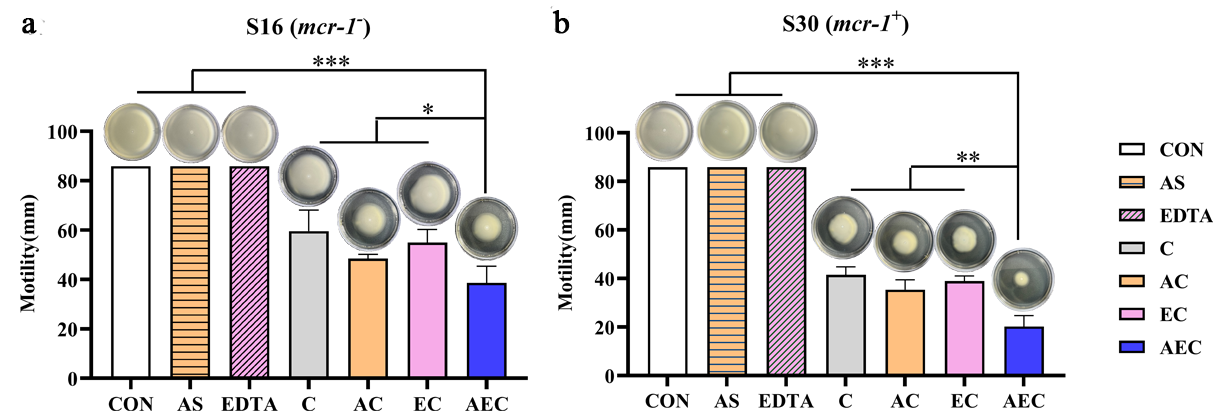

**Figure supplement 5. AS, EDTA, and COL inhibited the swimming motility of S16 (a) and S30 (b) strains.** Overnight cultures were were diluted 1:100 in fresh LB medium and grown to an OD_600_ of 0.5, and inoculated on 0.3% agar plates for 48 h at 37℃. The different concentrations of COL (2 mg/L) alone or in combination with 1/8 MIC of AS (156.3 mg/L) or EDTA (15.6 mg/L) were add into plates. The migration distance (cm) was measured. Data were shown in the mean of triplicates ± SD (*** *p* < 0.001, ** *p* < 0.01, * *p* < 0.05).

**Table supplement 1. The antibacterial activities of COL, AS and EDTA against the tested strains after single and double combinations**

| Strains | *mcr-1* | MICs (mg/L) | | | | | | | | | | |
| --- | --- | --- | --- | --- | --- | --- | --- | --- | --- | --- | --- | --- |
|  |  | Alone | | | COL + AS | | | | COL + EDTA | | | |
|  |  | COL | AS | EDTA | 1/4  AS | 1/8  AS | 1/16  AS | **Fold**  **change** | 1/4  EDTA | 1/8  EDTA | 1/16  EDTA | **Fold**  **change** |
| JS | － | 0.25 | 1250 | 125 | 0.0625 | 0.125 | 0.125 | **2-4** | 0.25 | 0.25 | 0.25 | **0** |
| S34 | － | 0.125 | 1250 | 125 | 0.008 | 0.015 | 0.125 | **0-16** | 0.015 | 0.03 | 0.0625 | **2-8** |
| S16 | － | 4 | 1250 | 125 | 0.25 | 0.25 | 4 | **0-****16** | 2 | 2 | 4 | **0-2** |
| S20 | － | 2 | 1250 | 125 | 0.015 | 0.015 | 2 | **0-133** | 0.25 | 2 | 2 | **0-8** |
| S13 | + | 2 | 1250 | 125 | 0.25 | 0.25 | 2 | **0-8** | 1 | 2 | 2 | **0-2** |
| S30 | + | 4 | 1250 | 125 | 0.25 | 0.25 | 2 | **2-16** | 2 | 2 | 2 | **2** |
| E16 | + | 2 | 1250 | 500 | 0.0625 | 0.25 | 2 | **0-32** | 0.5 | 0.5 | 1 | **2-4** |
| M15 | － | ＞64 | 1250 | ＞1000 | ＞64 | ＞64 | ＞64 | **0** | ＞64 | ＞64 | ＞64 | **0** |
| P01 | － | ＞64 | 1250 | ＞1000 | ＞64 | ＞64 | ＞64 | **0** | ＞64 | ＞64 | ＞64 | **0** |

**Table supplement 2. MIC values of COL against S16 and S30 strains after addition of exogenous** **cations**

| Strains | Cations  (100 mg/L) | MICs (mg/L) | | | |
| --- | --- | --- | --- | --- | --- |
|  |  | COL | AC | EC | AEC |
| S16  (*mcr*-*1*^-^) | Control | 4 | 0.25 | 2 | 0.125 |
|  | Na^+^ | 4 | 0.25 | 2 | 0.125 |
|  | K^+^ | 4 | 0.25 | 2 | 0.125 |
|  | Ca^2+^ | 4 | 0.25 | 2 | 0.125 |
|  | Mg^2+^ | 8 | 0.5 | 4 | 0.25 |
|  | Mn^2+^ | 4 | 0.25 | 2 | 0.125 |
|  | Zn^2+^ | 4 | 0.25 | 2 | 0.125 |
| S30  (*mcr*-*1*^+^) | Control | 4 | 0.25 | 2 | 0.125 |
|  | Na^+^ | 4 | 0.25 | 2 | 0.125 |
|  | K^+^ | 4 | 0.25 | 2 | 0.125 |
|  | Ca^2+^ | 4 | 0.25 | 2 | 0.125 |
|  | Mg^2+^ | 8 | 0.5 | 4 | 0.25 |
|  | Mn^2+^ | 4 | 0.25 | 2 | 0.125 |
|  | Zn^2+^ | 4 | 0.25 | 2 | 0.125 |

**Table supplement 3. MIC values of COL against S16 and S30 strains after addition of exogenous LPS**

| Strains | Drug | MICs (mg/L) | | | |
| --- | --- | --- | --- | --- | --- |
|  |  | + 0 LPS | + 4 LPS | + 32 LPS | + 512 LPS |
| S16  (*mcr*-*1*^-^) | COL | 4 | 4 | 8 | 64 |
|  | AC | 0.25 | 0.5 | 2 | 32 |
|  | EC | 2 | 2 | 2 | 32 |
|  | AEC | 0.125 | 0.5 | 2 | 32 |
| S30  (*mcr*-*1*^+^) | COL | 4 | 4 | 16 | 64 |
|  | AC | 0.25 | 0.5 | 4 | 64 |
|  | EC | 2 | 4 | 16 | 64 |
|  | AEC | 0.125 | 0.5 | 4 | 64 |

**Table supplement 4. MIC values of COL against S16 strain after the overexpression of different genes**

| Genes | L-Ara | MICs (mg/L) | | |
| --- | --- | --- | --- | --- |
|  |  | AC | EC | AEC |
| *cheA* | — | 0.25 | 0.5 | 0.125 |
|  | + | 0.25 | 2 | 4 |
| *cheY* | — | 0.25 | 2 | 0.125 |
|  | + | 0.5 | 2 | 0.125 |
| *STMDT2-34621* | — | 0.25 | 1 | 0.125 |
|  | + | 0.5 | 1 | 0.25 |
| *aer* | — | 0.25 | 1 | 0.125 |
|  | + | 0.25 | 1 | 0.125 |
| *fliD* | — | 0.25 | 1 | 0.25 |
|  | + | 0.5 | 1 | 0.5 |
| *fliT* | — | 0.25 | 1 | 0.125 |
|  | + | 0.25 | 1 | 0.125 |
| *opuBB* | — | 0.25 | 1 | 0.125 |
|  | + | 0.25 | 1 | 0.125 |
| *gltI* | — | 0.25 | 1 | 0.125 |
|  | + | 0.25 | 1 | 0.125 |
| *dppB* | — | 0.25 | 1 | 0.125 |
|  | + | 0.25 | 1 | 0.125 |
| *dppC* | — | 0.25 | 1 | 0.125 |
|  | + | 0.25 | 1 | 0.125 |
| *spvD* | — | 0.25 | 1 | 0.125 |
|  | + | 1 | 1 | 0.5 |

**Table supplement 5. MIC values of COL against S30 strain after the incubation of polypeptides**

| Strains | Drug | MICs (mg/L) | | |
| --- | --- | --- | --- | --- |
|  |  | Control | + P_u_ | + P_m_ |
| S30  (*mcr*-*1*^+^) | AC | 0.25 | 2 | 0.25 |
|  | AEC | 0.125 | 1 | 1 |

Note: The “P_u_” and “P_m_”indicate the unmutated and mutated peptide at THR 283, SER 284, and TYR 287 sties, respectively.

**Table supplement 6. Sequences of primers used in this study**

| **Primer** | **Sequence (5’ → 3’)** | **References** |
| --- | --- | --- |
| *cheA* – F | AATCTCGAGGTGAGCATGGATATTAGCGA | This study |
| *cheA* – R | AATGAATTCTCAGGCGGCTGTGATCGCCA |  |
| *cheY* – F | ACACTCGAGATGGCGGATAAAGAGCTTAA | This study |
| *cheY* – R | ACAGAATTCTCACATGCCCAGTTTCTCAA |  |
| *STMDT2_34621* - F | GGCCTCGAGATGAAAAATATCAAAGTCATCAC | This study |
| *STMDT2_34621* - R | GGCGAATTCTTAGAAGCTTTCCCAGTTCG |  |
| *aer* - F | AATCTCGAGATGTCTTCTCATCCCTACGT | This study |
| *aer* - R | GGCCTGCAGTTAATGCAGTACCGTGA |  |
| *fliD* - F | GGCCTCGAGATGGCTTCAATTTCATCATT | This study |
| *fliD* - R | AACCTGCAGTCAGGACTTGTTCATAGCT |  |
| *fliT* - F | AATCTCGAGATGACCTCAACCGTGGA | This study |
| *fliT* - R | AATGAATTCTTATGAGGCGCCAGGCG |  |
| *oppuBB* - F | GGCCTCGAGATGGATACGATACATTATATG | This study |
| *oppuBB* - R | AATGAATTCTTATCGTATCCCCTTCGGTG |  |
| *gltI* - F | GGCCTCGAGATGATAAAGGAGTTGGATAT | This study |
| *gltI* - R | GGCGAATTCTTAGTTAAGCGCTTTATCATTC |  |
| *dppB* - F | AATCTCGAGATGTTGCAGTTCATTCTCCG | This study |
| *dppB* - R | GGCGAATTCTTACTTCTTATGCCGAATACGC |  |
| *dppC* - F | AGGCTCGAGATGTCACAGGTTACTGAAAA | This study |
| *dppC* - R | AGGGAATTCTTACTGCTTCAGTTTGGGATC |  |
| *spvD* - F | AATCTCGAGATGAGAGTTTCTGGTAGTGC | This study |
| *spvD* - R | GGCGAATTCTCAATCGTGTTTTTCATCAT |  |
| *mcr*-*1* - qF | TGCTCCAAAATGCCCTACAGACC | Yi *et al*, 2022 |
| *mcr*-*1* - qR | TGCCCCAAGTCGGATAATCCAC |  |
| 16S rRNA - qF | TGTCGTCAGCTCGTGTTGTG | Yi *et al*, 2022 |
| 16S rRNA - qR | ATCCCCACCTTCCTCCAGTT |  |

**Table supplement 7. The antibacterial activities of COL, AS and EDTA against the tested strains after single and double combinations**

| Strains | *mcr-1* | MICs (mg/L) | | | | | | | | | | |
| --- | --- | --- | --- | --- | --- | --- | --- | --- | --- | --- | --- | --- |
|  |  | Alone | | | COL + AS | | | | COL + EDTA | | | |
|  |  | COL | AS | EDTA | 1/4  AS | 1/8  AS | 1/16  AS | **Fold**  **change** | 1/4  EDTA | 1/8  EDTA | 1/16  EDTA | **Fold**  **change** |
| S29 | + | 2 | 1250 | 125 | 0.015 | 0.125 | 0.5 | **4-133** | 1 | 1 | 2 | **0-2** |
| S31 | + | 4 | 1250 | 125 | 0.25 | 0.5 | 2 | **2-16** | 2 | 2 | 2 | **0** |
| S23 | + | 4 | 1250 | 125 | 0.25 | 0.5 | 1 | **4-16** | 2 | 2 | 2 | **0** |
| S93 | + | 4 | 1250 | 125 | 0.25 | 2 | 2 | **2-16** | 2 | 2 | 2 | **0** |

**Table supplement 8. The antibacterial activities of COL against the tested strains after single and triple combinations**

| Strains | MICs (mg/L) | | | | | | | | | **Fold change** |
| --- | --- | --- | --- | --- | --- | --- | --- | --- | --- | --- |
|  | COL  alone | COL + 1/4 AS +EDTA | | | | COL + 1/8 AS +EDTA | | | |  |
|  |  | 1/4  EDTA | 1/8  EDTA | 1/16  EDTA | 1/32  EDTA | 1/4  EDTA | 1/8  EDTA | 1/16  EDTA | 1/23  EDTA |  |
| S29 | 2 | 0.00003 | 0.001 | 0.03 | 0.0625 | 0.008 | 0.03 | 0.125 | 0.125 | **16-66667** |
| S31 | 4 | 0.03 | 0.25 | 0.25 | 0.5 | 0.125 | 0.25 | 0.5 | 0.5 | **8-133** |
| S23 | 4 | 0.0625 | 0.125 | 0.125 | 0.25 | 0.125 | 0.25 | 0.5 | 0.5 | **8-64** |
| S93 | 4 | 0.03 | 0.0625 | 0.125 | 0.25 | 0.25 | 0.5 | 0.5 | 2 | **2-133** |

**Table supplement 9. The MICs of different antimicrobial drugs against the S16 and S30**

| Drug | FFC | ENR | DOX | AMP | AMK | FOS | CAZ | CRO | CTX | TGC |
| --- | --- | --- | --- | --- | --- | --- | --- | --- | --- | --- |
| Breakpoint | ≥16 | 2 | ≥16 | ≥32 | ≥16 | ≥256 | ≥16 | ≥4 | ≥4 | ≥4 |
| S16 | 2 | ＜0.5 | 2 | 512 | ＜0.5 | 2 | ＜0.5 | ＜0.5 | ＜0.5 | 4 |
| S30 | 128 | 2 | 32 | ＞512 | 1 | 8 | ＜0.5 | ＜0.5 | ＜0.5 | 4 |

Note: FFC: Flufenicol; ENR: Enrofloxacin; DOX: Doxycycline; AMP: Ampicillin; AMK: Amikacin; FOX: Fosfomycin; CAZ: ceftazidime; CRO: Ceftriaxone; CTX:Cefotaxime; TGC: Tigecycline.

**Supplement materials and methods**

**Overexpression of SDEGs in S16 Strain**

The complete open reading frame of SDEGs (including *cheA*, *cheY*, *STMDT2-34621*, *aer*, *fliD*, *fliT*, *opuBB*, *gltI*, *dppB*, and *spvD*) were amplified by PCR from the genomic DNA of strain S16. Then these genes were inserted into the multiple cloning site of vector pBAD (Ampicillin^+^), and the recombinant plasmids were chemically transformed into S16 cells. Finally, the target genes were induced by 0.2% L-arabinose (L-Ara) when necessary.

**RT-PCR analysis**

The *mcr*-*1*^+^ *Salmonella* S30 strain was grown to OD_600_ ≈ 0.5, then incubated with double and triple combination strategies (same as the Time-kill assays section) for 6 h. Total RNA were extracted isolated by following the phenol-chloroform method, and then were reverse transcribed to cDNA using PrimeScript™RT reagent Kit (TaKaRa). Primers for RT-PCR were designed according to (Yi *et al*, 2022), listed in Table supplement 6. The 16S rRNA gene was chosen as a housekeeping gene. The relative expression ratio of the gene tested were determined using 2^−∆ (∆CT)^ method, as compared to that of AS and EDTA treatment.

**Molecular docking**

Crystal structure of the C-terminal catalytic domain of MCR-1 with two Zinc ions (PDB ID: 5GRR) was obtained from the Protein Data Bank (PDB, http://www.rcsb.org). 3D structure of AS was downloaded from The Pubchem Project (PubChem CID: 6917864). The region containing the previously reported active site within MCR-1 was defined as the binding site for docking simulations. AutoDock software was used for the flexible ligand docking between MCR-1 and AS. The results that were ranked permitting to energy values of the docking (kcal/mol), and the lower the value, the more likely the ligand-active site bind. The Discovery Studio molecular graphics system was used to preliminarily estimate and further confirm the modes of interaction with binding site residues.

**Competitive inhibitory assays**

Two polypeptides from MCR-1 (named as P_u_ and P_m_) were synthesized by Sangon Biotech Company. P_u_ (5’-CG**TS**TA**Y**SVP-3’) and P_m_ (5’-CG**AA**TA**A**SVP-3’) were unmutated and mutated peptides from the binding sites THR283, SER284, and TYR287, respectively. AS and AS+EDTA were pre-incubated with P_u_ or P_m_ for 2 h, and then added into the *mcr*-*1*^+^ *Salmonella* S30 strain. The MIC of COL were determined by the 2-fold serial broth microdilution method according to CLSI guidelines (Wayne, 2021).

**Motlity assays**

Motlity assays were performed using 0.3% agar plates containing AS, EDTA, COL alone or drug combinations. The final concentration of COL was 2 mg/L, when used alone or in drug combinations. AS and EDTA were added at final concentrations equivalent to their 1/8 MICs, when used in drug combinations. Overnight culture of S16 (*mcr*-*1^-^*) and S30 (*mcr*-*1*^+^) was diluted 1:100 in fresh LB medium and grown to an OD_600_ of 0.5, and inoculated on 0.3% agar plates. The migration distance (cm) was measured and recorded for 48 h at 37 ℃.
